# A Full-Thickness 3D Bioprinted Melanoma-on-Chip Platform for Preclinical Therapeutic Testing

**DOI:** 10.64898/2026.11.23.768514

**Authors:** Alexeja Kleiter, Nora Kaiser, Elena Brunner, Andreas Aufschnaiter, Ema Husarčíková, Guido Wollmann, Beate Rinner, Daniela Anna Ortner-Tobider, Patrizia Stoitzner, Judith Hagenbuchner, Michael J. Ausserlechner

## Abstract

Malignant melanoma remains a major clinical challenge due to its high metastatic potential and frequent therapy resistance. Although recent therapeutic advances have improved patient survival, predictive preclinical models that recapitulate the human tumor microenvironment (TME) remain limited. We developed a human 3D bioprinted melanoma-on-chip platform that combines a fully differentiated, barrier-forming human skin model with microfluidic perfusion and spatially controlled integration of melanoma spheroids at the epidermal-dermal junction. This provides a standardized tissue environment for longitudinal tumor and therapeutic studies for up to 21 days. To validate our model for preclinical therapeutic testing, we treated the melanoma-on-chip with the oncolytic virus VSV-GP. In 2D, viral infection of melanoma cells resulted in homogeneous cell death within 48 hours. In melanoma-on-chip, viral propagation was restricted to tumor cells and markedly delayed compared to 2D. Although VSV-GP stopped melanoma growth in 3D, it did not achieve complete tumor-cell eradication, as observed in 2D culture. These findings demonstrate that the 3D skin tissue environment substantially influences viral infection kinetics and therapeutic outcome, underscoring the need for more physiologically relevant human tissue models in preclinical drug evaluation.

## Introduction

Melanoma accounts for the majority of skin cancer-related deaths despite representing only a small proportion of all cutaneous malignancies [1–3]. The disease arises from malignant transformation of melanocytes, where oncogenic mutations drive uncontrolled proliferation and metastasis [2]. While current treatments have improved patient outcomes, therapeutic resistance and disease recurrence remain critical clinical challenges [4]. Recent advances in immune checkpoint inhibition have fundamentally reshaped the treatment landscape of metastatic melanoma. However, a substantial number of patients still exhibit primary resistance or develop acquired resistance after an initial response [5, 6]. In metastatic melanoma, accumulating evidence suggests that these responses are not driven solely by tumor-intrinsic factors but are also strongly influenced by the local TME [7, 8]. Conventional 2D cell cultures provide simple conditions but fail to recapitulate the architecture, mechanical properties, tissue stiffness, extracellular matrix composition, and multicellular interactions that shape tumor behavior. Recent advances in melanoma biofabrication have addressed several of these limitations by incorporating multiple cell types, extracellular matrix components, spatial control, and microfluidic culture into increasingly complex human tissue models [9–12]. However, existing approaches generally capture only selected aspects of the melanoma microenvironment, leaving the need for biofabricated melanoma models that combine anatomical tissue organization with reproducible fabrication, long-term culture, and functional therapeutic testing within a single tissue construct. [13–15].

3D bioprinting-on-chip offers an effective way of assembling tissue equivalents directly within fluidic chip devices. By integrating laser-manufactured fluidic chips with extrusion- and microjet-based bioprinting, viable tissue can be directly fabricated within customizable microfluidic chip devices using different cell-laden bioinks. This enables the spatially controlled integration of multiple cell types and biomaterials to generate multilayered and multicellular tissue. The 3D bioprinted tissue can be conveniently handled, perfused, maintained in long-term culture, imaged, and functionally analyzed. These advanced biofabricated tissue models also provide a platform for investigating the efficacy of emerging cancer therapies, such as oncolytic viruses (OV) [14–17].

OV preferentially infect and replicate in malignant cells, kill tumor cells and ultimately lead to an immune cell influx and anti-tumor immune response. In this project, we used the negative-strand RNA oncolytic virus VSV-GP [18]. This virus is a chimeric vesicular stomatitis virus encoding the glycoprotein of the lymphocytic choriomeningitis virus and has demonstrated promising anti-tumor activity and an improved safety profile compared to wild-type VSV [18–21]. The efficacy of OV is greatly influenced by the TME, which includes tissue architecture, extracellular matrix composition, stromal interactions, and antiviral signaling. In 2D, unrestricted viral access results in homogeneous infection and rapid replication, leading to an overestimation of therapeutic efficacy under physiologically simplified conditions. Species-specific differences in antiviral signaling, complex interactions of TME and tumor cells and other factors heavily influence the resulting replication kinetics, limiting interpretability of simple mouse or human 2D models [18, 22–24]. Therefore, human 3D tissue models are valuable intermediate platforms for investigating virus-tumor interactions and therapeutic efficacy under standardized conditions that more accurately reflect the human melanoma microenvironment.

In this paper, we describe a human melanoma-on-chip model that recapitulates key structural features of the human melanoma tissue environment. It integrates a fully differentiated epidermis, spatially controlled melanoma positioning at the epidermal-dermal junction, microfluidic perfusion inside the multilayered dermis, long-term tissue culture, and direct therapeutic assessment within a single biofabricated construct. As proof of concept, we also demonstrate the model’s applicability in evaluating therapeutic responses to anti-tumor therapy. We hypothesized that embedding melanoma within a reconstructed human skin environment would alter viral replication kinetics and therapeutic efficacy compared to conventional 2D culture.

## Results

### 3D bioprinted skin in laser-manufactured microfluidic PMMA chips

To replicate human skin physiology, we developed a custom-designed microfluidic chip platform that supports dermal and epidermal maturation under controlled culture conditions. Therefore, we designed laser-fabricated microfluidic chips made of polymethyl methacrylate (PMMA) for perfused and static culture. The chip enabled long-term tissue culture with continuous nutrient supply and direct access to the anchored skin tissue. PMMA was chosen for its non-toxicity, optical transparency, mechanical stability, ease of laser processing, and compatibility with rapid, cost-effective fabrication of a customized chip. Using an industrial laser engraving system, chips were manufactured with a central tissue chamber connected to perfusion channels for continuous medium exchange and, in the static setting, four additional orthogonal channels provided supplementary medium supply when these chips were cultured in 6-well plates (Figure 1a). The laser mode and pulse frequency were adjusted to specifically roughen the PMMA surface for enhanced tissue adhesion. The PMMA chip was permanently bonded to a glass coverslip using a biocompatible double-sided adhesive, yielding robust, reusable chip assemblies suitable for prolonged culture. Prior to the bioprinting process, the chip was treated with oxygen plasma to activate the surface for improved biomatrix adhesion and wetting.

**Figure 1.**
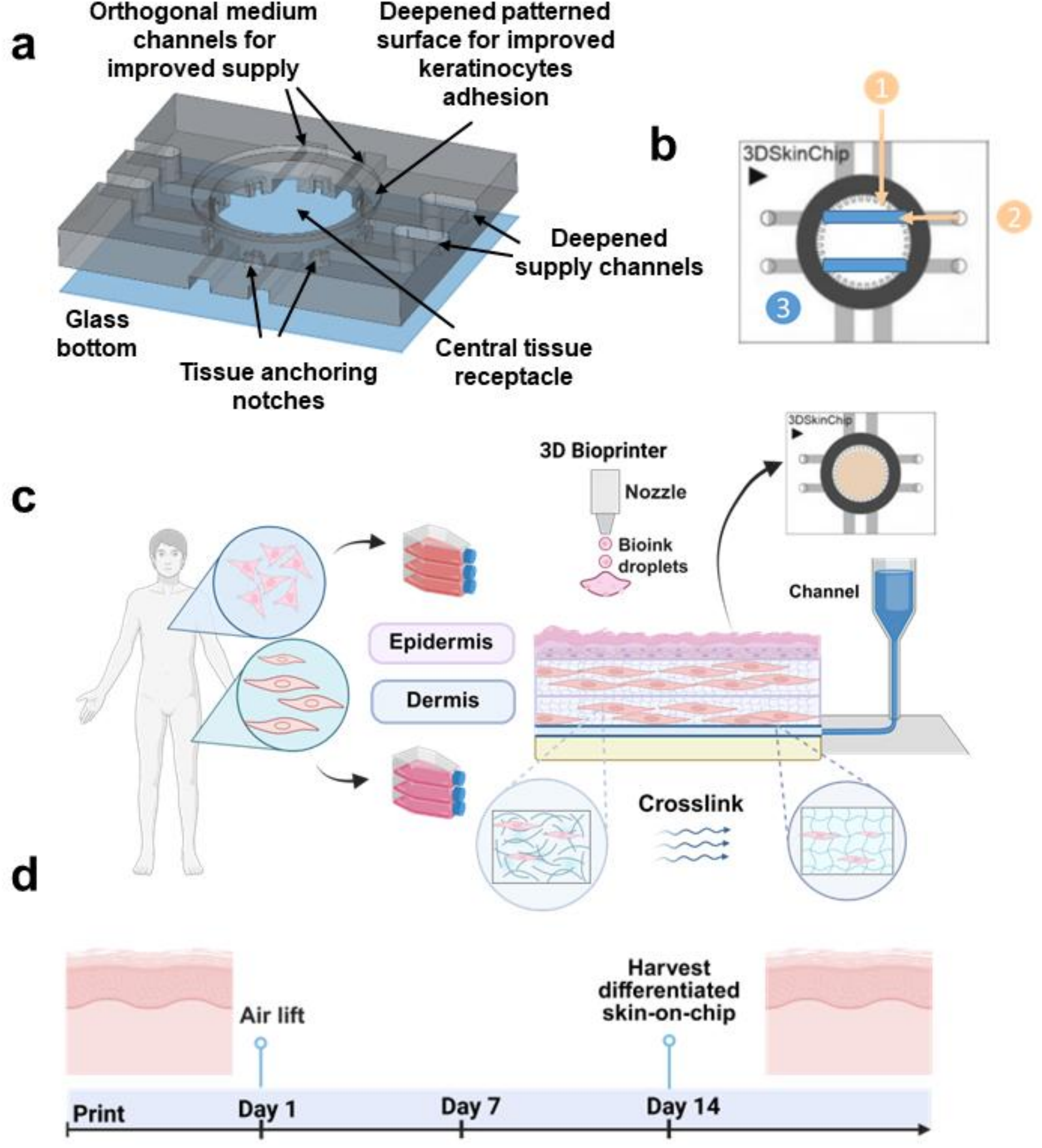
Fluidic chip devices for 3D bioprinted skin-on-chip. (a) CAD designs of the microfluidic PMMA chips for skin culture with detailed description of features. (b) Top view schematic of the assembled PMMA chip (3DSkinChip). B1: orthogonal microchannels for additional supply of the central tissue chamber. B2: engraved microfluidic channels within the chip. B3: Pluronic F127 templated microchannels within the dermal compartment directly connected with engraved channels for medium perfusion. (c) Schematic overview of the 3D bioprinting workflow for the skin-on-chip model. Dermal fibroblasts printed as sequential bioink layers (dermis with channels followed by epidermis) into chip device, followed by photocrosslinking. Keratinocyte layer microjetted on top using PVP. (d) Timeline of skin-on-chip fabrication and culture. Skin-on-chip bioprinted on d0 and transferred to an air-liquid interface on d1. Differentiated skin harvested on d14 (Schematics created with BioRender).

Skin-on-chip was fabricated directly within the microfluidic chambers using a RegenHU 3D Discovery BioSafety bioprinter equipped with extrusion and micro-jetting printheads (Supplementary Figure S1a). To generate hollow channels within the dermal compartment, microchannel structures were printed using Pluronic F127 as thermo-reversible sacrificial ink (Figure 1b and Supplementary Figure S1b). Following deposition and photo-crosslinking of the surrounding hydrogel, dissolution of the Pluronic strands generated hollow, interconnected channels that enabled continuous medium flow throughout the construct. The dermal compartment was generated by micro-jetting of human dermal fibroblasts (HDFs) suspended in a gelatin methacryloyl (GelMA)-based hydrogel supplemented with 10% collagen and lithium phenyl (2,4,6-trimethylbenzoyl) phosphinate (LAP). Printing was performed using a heated printhead to deposit three sequential hydrogel layers, each stabilized by 10 seconds of photo-crosslinking (Supplementary Figure S1b). Keratinocytes were suspended in a polyvinylpyrrolidone (PVP) and medium mix-based formulation to ensure homogeneous cell distribution on top of the 3D bioprinted tissue (Figure 1c). The cells were printed as a single epidermal layer using a heated printhead. This workflow enabled the fabrication of 12 skin-on-chip models with an average printing time of 2 minutes per skin-chip. Following fabrication, constructs were maintained under submerged conditions for 24 h before keratinocytes were exposed to air (air-liquid interface, ALI) and further cultured in keratinocyte differentiation medium for 14 days (Figure 1d). The deepened supply channels allowed medium delivery, while the central chamber remained uncovered in static culture in 6-well plates. To further investigate the impact of chip design on tissue development, two alternative stereolithography (SLA)-printed chip prototypes were evaluated. Both designs incorporated a supporting grid structure beneath the dermal compartment and differed in the presence or absence of a Pluronic-based channel in the center of the dermis (Supplementary Figure S2a). Although Chip 2 and Chip 3 prototypes ensured larger medium reservoirs, they consistently led to reduced fibroblast viability as indicated by loss of fluorescence intensity (Supplementary Figure S2b). Histological analysis revealed malformed epidermal tissue architecture in Chip 2 and Chip 3. Furthermore, vimentin staining showed an absence of elongated fibroblasts after 14 days of culture, indicating impaired fibroblast morphology under these culture conditions (Supplementary Figure S2c). Based on these observations, neither design was considered suitable for maintaining the desired tissue characteristics during long-term culture.

### Optimization of the 3D bioprinted dermis

Successful engineering of human skin equivalents requires a dermal matrix which provides sufficient mechanical stability while supporting fibroblast viability and stretching. To identify the optimal hydrogel composition, GelMA concentrations of 5% and 7.5%, each supplemented with 10% collagen, were evaluated for their mechanical and biological performance. Rheological characterization revealed a concentration-dependent increase in hydrogel stiffness: the shear storage modulus (G′) increased from approximately 3 kPa for 5% GelMA to 9 to 12 kPa for 7.5% GelMA. This confirms that increasing GelMA concentration substantially improved the stiffness of the bioink (Figure 2a), following a non-linear relation between concentration and stiffness when analyzing the shear storage modulus of 3%, 4%, 5%, 7.5% and 10% GelMA (Supplementary Figure S3). Histological analysis revealed that constructs generated with 7.5% GelMA closely resembled human skin, with a dermal thickness of approximately 1.3 mm (Figure 2b). Both hydrogel formulations supported fibroblast survival and stretching within the printed dermal compartment. Vimentin staining confirmed the mesenchymal phenotype of HDFs and their elongated morphology after two weeks in culture in 5% GelMA and 7.5% GelMA conditions, as opposed to 10 % GelMA (Figure 2c). This condition was excluded from further experiments, as its high stiffness restricted fibroblasts from adopting their characteristic elongated morphology. Fluorescence microscopy of EYFP-expressing HDFs demonstrated stable EYFP expression throughout the culture period. After detaching the fibroblasts from the cell culture flask and after printing, HDFs displayed a predominantly rounded morphology that progressively transitioned into the characteristic spindle-shaped phenotype found in human skin (Figure 2d). Quantitative image analysis confirmed this transition, with the circularity index significantly decreasing from 0.45 on d1 to 0.35 after d14, indicating progressive cell elongation and adaptation within the GelMA and collagen matrix quantified using ImageJ (Figure 2e). We compared the optimized PMMA-based microfluidic platform with two previously developed SLA-printed chips made of biocompatible resin. Although both systems supported initial tissue formation, the SLA-based chips showed a progressive loss of EYFP fluorescence, indicating reduced fibroblast viability, together with impaired keratinocyte differentiation and altered dermal and epidermal thickness over prolonged culture (Supplementary Figure S2b and c). These findings demonstrate that combining the optimized 7.5% GelMA and collagen bioink with the laser-manufactured PMMA chip provides improved dermal thickness, stability and supports reproducible, long-term generation of human dermal tissue.

**Figure 2.**
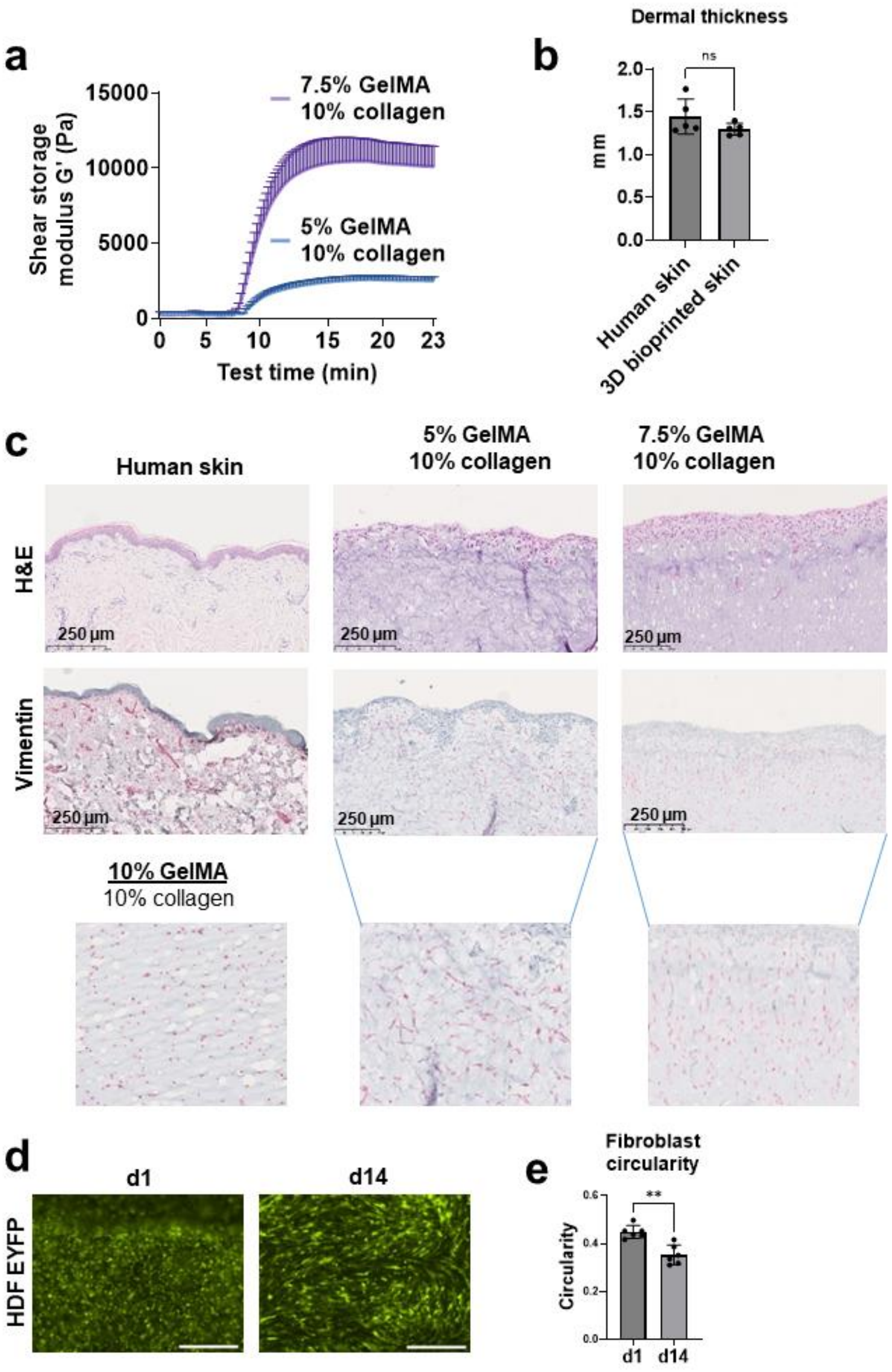
Optimization of the 3D bioprinted dermal compartment. (a) Rheological analysis of mechanical stiffness (storage modulus G’) for 5% GelMA and 7.5% GelMA hydrogels containing 0.25% LAP, during crosslinking with violet light (405 nm). (b) Dermal thickness measured from H&E-stained histological sections of human skin and 3D bioprinted skin constructs cultured in vitro. Data are presented as mean ± SD. Statistical analysis was performed using an unpaired t-test (P = 0.1605). (c) Histological characterization of the 3D bioprinted dermis after 2 weeks of culture, compared with human skin. H&E (top) and vimentin (middle) staining of human skin (left), dermal constructs generated with 5% GelMA plus 10% collagen (middle), and 7.5% GelMA plus 10% collagen (right). An additional condition of 10% GelMA plus 10% collagen (bottom) shows rounded fibroblasts. Scale bar 250 µm; 10× magnification. (d) Fluorescence microscopy of HDF/hTERT-EYFP fibroblasts on d1 and d14 in culture. Scale 500 µm; 20× magnification. (e) Fibroblast circularity index at d1 and d14 in constructs containing 7.5% GelMA and 10% collagen. Circularity was quantified using ImageJ. Data are presented as mean ± SD; statistical analysis was performed using Welch’s t-test (**P = 0.0012).

### Optimized epidermal differentiation and maturation of 3D bioprinted skin

Establishing a functional epidermal compartment requires highly regulated keratinocyte proliferation and differentiation. Therefore, bioprinted skin equivalents were cultured under ALI conditions for 14 days in keratinocyte differentiation medium. To investigate the influence of dermal stiffness on epidermal development, keratinocytes were printed onto dermal equivalents composed of either 5% or 7.5% GelMA supplemented with 10% collagen. Histological analysis revealed marked differences between the formulations. Constructs containing 5% GelMA/collagen developed a considerably thinner epidermal layer despite comparable fibroblast morphology, with keratinocyte islets also growing in the dermal layer and no clear separation between dermis and epidermis (Figure 3a top images). In human skin, proliferative activity in the epidermis is largely confined to the basal layer. In 5% and 7.5% GelMA/collagen dermal bioinks, Ki67-positive, proliferating keratinocytes were not confined to the boundary between dermis and epidermis as in human skin (Figure 3a, lower images) but were visible throughout the epidermis. To reduce humidity at the ALI, we added a daily drying step for 30 min exposure to airflow in a sterile hood. This controlled dehydration step markedly improved epidermal differentiation and restricted proliferating cells to the basal layer, thereby more closely resembling the spatial organization of human epidermis (Figure 3a, right image). Under these optimized conditions, epidermal thickness reached approximately 80–110 µm, closely resembling the architecture and marker expression in human epidermis (Figure 3b). Constructs maintained under ALI conditions developed a continuous, multilayered epidermis expressing the differentiation markers cytokeratin 1 (CK1) and cytokeratin 10 (CK10), whereas submerged cultures failed to express CK1/10, as demonstrated by automated capillary Western blot analysis (Figure 3c). The daily drying step showed significantly increased expression of the epidermal differentiation markers CK1/10 compared to the ALI condition (Figure 3d). In addition to CK1/10, involucrin was also detected by immunofluorescence staining (Figure 3e). To evaluate whether the engineered epidermis developed a functional barrier, a Lucifer Yellow permeability assay was performed after two weeks of culture. Under optimized culture conditions, only minimal dye penetration was observed, indicating the formation of a highly intact epidermal barrier. When the epidermal barrier was damaged by pre-incubation with sodium dodecyl sulfate (SDS), pronounced Lucifer Yellow penetration throughout the tissue and into the culture medium was observed, demonstrating a compromised barrier function (Figure 3f). Thus, a 7.5% GelMA/collagen dermal matrix combined with daily extra drying steps promoted the formation of reproducible and physiologically relevant whole skin equivalents.

**Figure 3.**
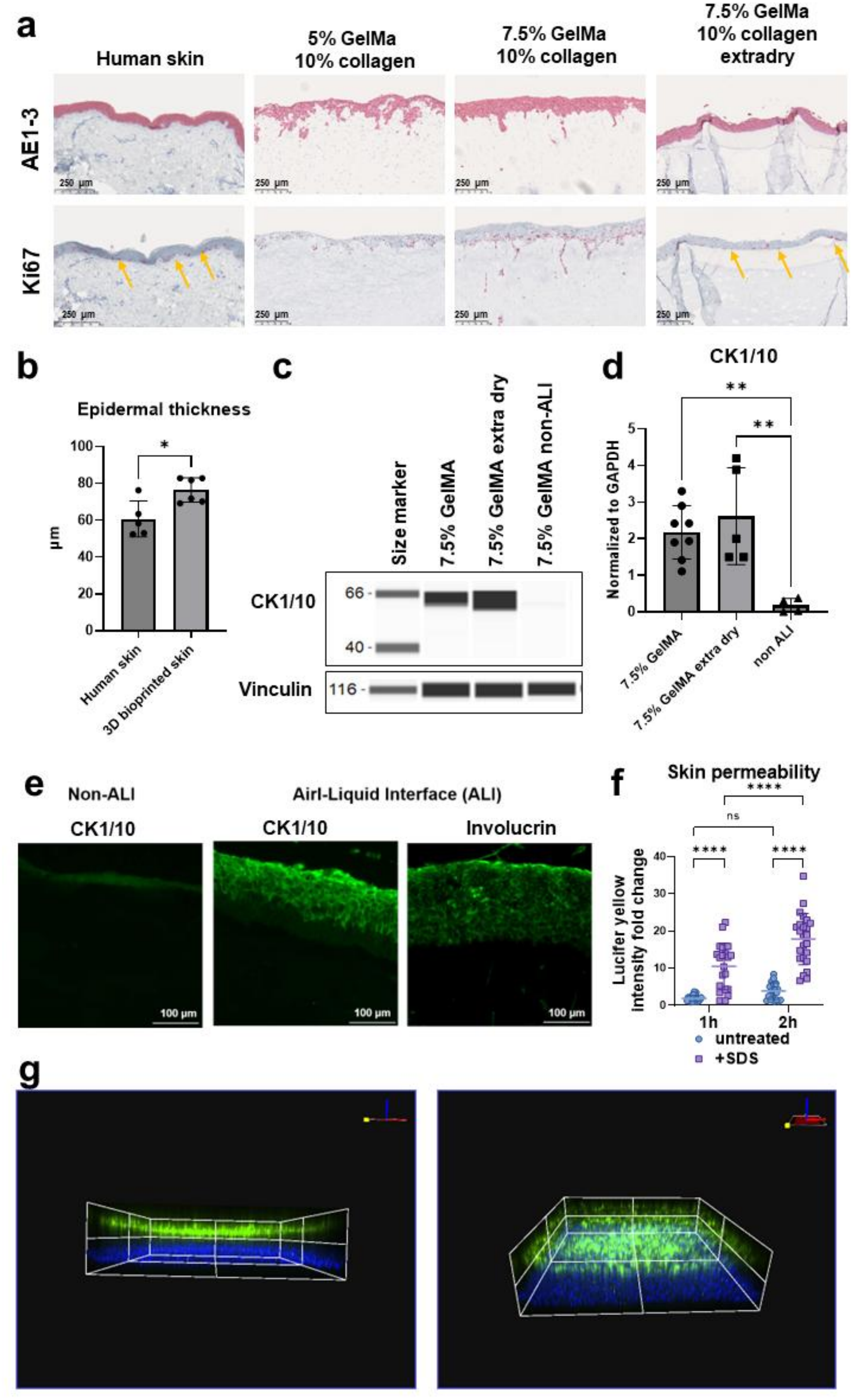
Epidermal differentiation and maturation in bioprinted skin equivalents. (a) Immunohistochemical characterization of 3D bioprinted skin cultured for 2 weeks, compared with human skin. AE1-3 (pan-cytokeratin) and Ki67 staining of human skin (left), 5% GelMA plus 10% collagen, 7.5% GelMA plus 10% collagen, and 7.5% GelMA plus 10% collagen with extra drying step (right). Arrows indicate Ki67-positive cells localized in close proximity to the basement membrane. Scale bar 250 µm; 10x magnification.(b) Epidermal thickness measured from H&E-stained histological sections of human skin and 3D bioprinted skin cultured in vitro. Data presented as mean ± SD. Statistical analysis was performed using the Mann-Whitney test (*P = 0.0303). (c) Automated capillary Western blot of CK1/10 expression, with vinculin as loading control. Lanes: 7.5% GelMA plus 10% collagen, equivalent constructs with extra drying step, 5% GelMA plus 10% collagen, and non air lifted (non ALI) control. (d) Quantification of CK1/10 expression normalized to GAPDH in 7.5% GelMA/10% collagen constructs, corresponding constructs subjected to additional drying, and non-ALI controls. Data are presented as mean ± SD. Statistical analysis was performed using two-way ANOVA (**P = 0.0043 (7.5% GelMA vs non ALI) and **P = 0.0019 (7.5% GelMA extra dry vs non ALI)). (e) Immunofluorescence staining of CK1/10 and involucrin after 2 weeks of culture. Comparison of non-ALI (top) and ALI-incubation. Scale bar is 100 µm. (f) Barrier function of bioprinted skin equivalents assessed by topical application of Lucifer Yellow, with or without 1% SDS pretreatment as a positive control for impaired skin integrity. Fold change in Lucifer Yellow fluorescence intensity in the medium reservoir was determined after 1 and 2 h. Data are presented as mean ± SD. Statistical analysis was performed using two-way ANOVA (ns = not significant= 0.1730 and **** P < 0.0001). (g) Representative 3D bioprinted skin-on-chip confocal images are shown from two viewing angles on d7 of ALI. The dermis (emiRFP670) is shown in blue, and the green-fluorescent epidermis (EYFP) is shown in green.

To visualize the distribution of different cell types also in living 3D bioprinted skin we expressed the near-infrared shifted protein emiRFP670 [25] in HDF and EYFP in keratinocytes. Although EYFP fluorescence partly diminishes during epidermal differentiation, live cell imaging enables 3D monitoring of skin constructs (Figure 3g). To create a vascularized full-thickness skin model, we coated the inner walls of the channels with human umbilical vein endothelial cells (HUVECs) using a specifically developed 3D shaker and coating protocol, as previously described [16]. Supplementary Figure S4 shows a 3D reconstruction of multi-color confocal image stacks through the vascularized skin equivalent. The entire wall of the channel was confluently lined with endothelial cells.

### Development of a patient-derived melanoma-on-chip model

To establish a physiologically relevant melanoma-on-chip model, we compared the 3D growth characteristics of the established melanoma cell line A375 and the patient-derived melanoma cells MUG-Mel2 and MUG-Mel3 (Supplementary Figure S5a). All three melanoma cell lines successfully formed spheroids, which were subsequently integrated into the 3D bioprinted skin-on-chip model (Supplementary Figure S5b). Since the A375 cell line carries numerous mutations, is hypotriploid, and has been in culture since 1973, we decided to use the recently isolated patient-derived MUG-Mel2 cells [26] for all subsequent experiments. An additional advantage of these melanoma cells is their stable melanin pigmentation that allows for label-free visualization and longitudinal monitoring of spheroid growth by brightfield microscopy. For the biofabrication workflow, preformed MUG-Mel2 spheroids were manually inserted into the still liquid microjet-printed top of the dermal fibroblast layer and then crosslinked by exposure to 405 nm light. Thereby anchoring the spheroids at the boundary between dermis and epidermis. The epidermal compartment was then microjet-printed on top (Figure 4a). The spatial positioning of the melanoma spheroids within the 3D bioprinted melanoma-on-chip proved to be critical. When the workflow included the different melanoma spheroids after the skin-on-chip print, spheroids grew on top of the keratinocyte layer (Supplementary Figure S5c). Melanoma-on-chips lacking keratinocytes exhibited a significant increase in spheroid growth normalized to day 1 (Supplementary Figure S5d). Macroscopic imaging the in Figure 4a described workflow demonstrated uniform incorporation of pigmented melanoma spheroids with an initial area of approximately 0.6 mm2 (Figure 4b left). Macroscopic imaging allowed for non-destructive monitoring and revealed a significant increase in tumor size over three weeks of culture at ALI, while maintaining a defined spatial position within the tissue construct. After 21 days in culture, the spheroids had an average area of 2.58 mm2 (Figure 4b and Figure 4c). Two days post printing, confocal imaging and fluorescence microscopy confirmed successful integration of the melanoma spheroid above the fibroblast dermal layer and in between the keratinocyte layer, thereby reflecting the anatomical position of human melanoma (Figure 4d and Supplementary Figure S5e). Hematoxylin and eosin (H&E) staining of paraffin-embedded melanoma-on-chip tissue after 21 days in culture also revealed that the melanoma spheroids remained localized at the epidermal-dermal junction. CK staining confirmed preservation of the epidermal compartment bordering the tumor, while S100 staining specifically identified melanoma cells within the tumor core. Ki67 staining visualized proliferative activity in both melanoma cells and basal keratinocytes (Figure 4e). Collectively, these findings demonstrate that the optimized bioprinting workflow enables reproducible integration of melanoma spheroids into engineered human skin while preserving tissue integrity, supporting long-term tumor growth, and allowing non-destructive longitudinal monitoring. The model therefore provides the basis for investigating melanoma progression and evaluating therapeutic responses within a controlled 3D human skin tissue environment.

**Figure 4.**
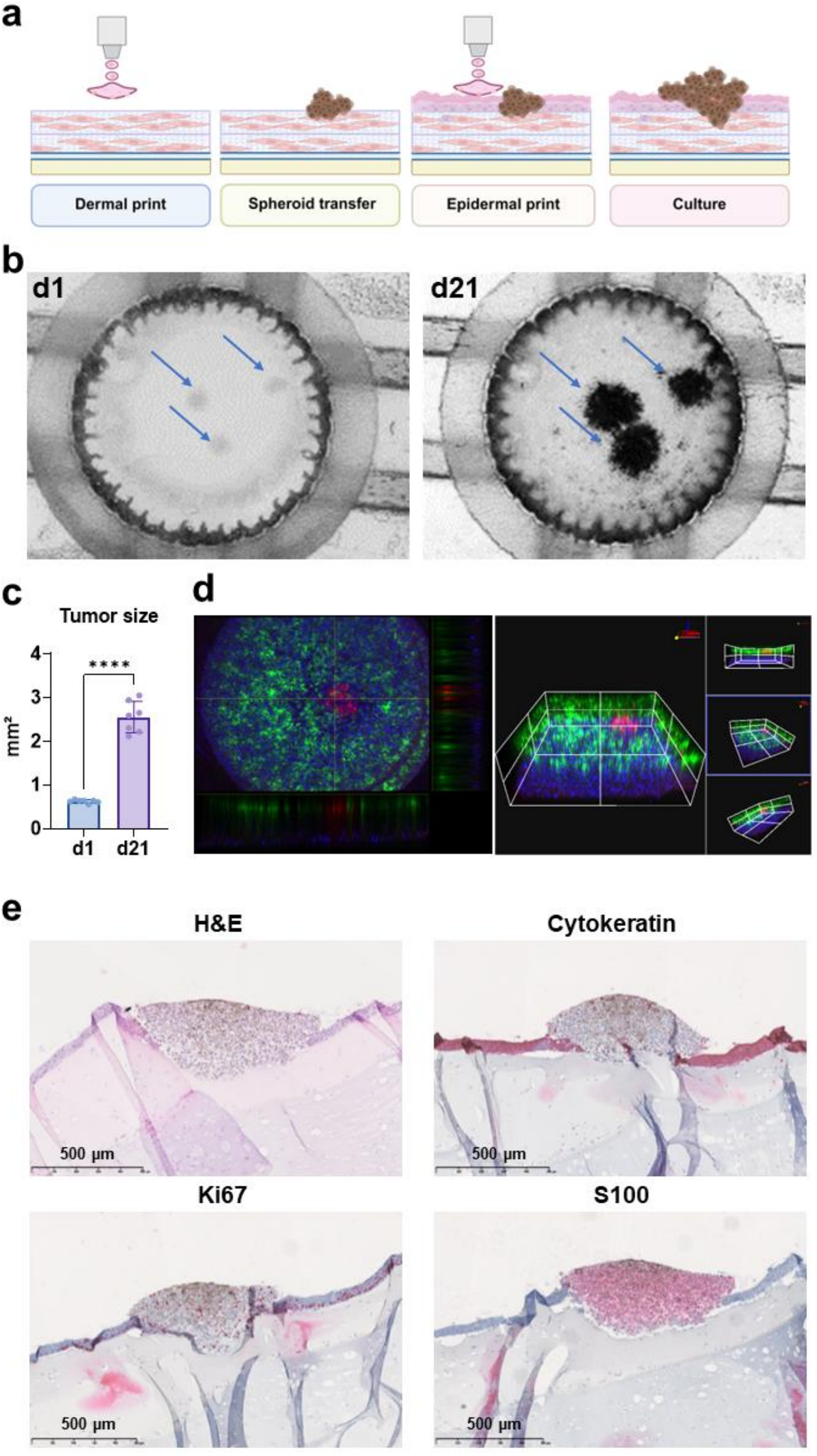
3D bioprinted melanoma-on-chip model. (a) Schematic overview of bioprinting procedure: dermal print, spheroid transfer, crosslinking with violet light, epidermal print, and culture at air-liquid interface. (b) Macroscopic top view images of melanoma-on-chip devices at d1 and d21, showing progressive darkening, growth and spreading of the melanoma tumors in skin-on-chip (arrows). Diameter of tissue receptacle is 1 cm. (c) Melanoma tumor size at d1 and d21. Data are mean ± SD; statistical significance was assessed using Welch’s t-test; n = 7 (**** P <0,0001). (d) Confocal imaging of a melanoma co-culture model. MUG-Mel2 melanoma cells expressing turboRFP (red), keratinocytes expressing EYFP (green), and HDFs expressing emiRFP670 (blue) were imaged using an FV4000 (Evident) confocal microscope (4× objective). (e) Immunohistochemical characterization of MUG-Mel2 tumors grown for three weeks in skin-on-chip: H&E, pan-cytokeratin, Ki67, and S100 staining. Scale bar 500 µm.

### VSV-GP-ZsGreen induces tumor-cell restricted destruction of the melanoma-on-chip

We next assessed the melanoma-on-chip model for preclinical therapeutic testing. We infected the melanoma-on-chip model grown for 7 days with a fluorescent reporter version of the oncolytic virus VSV-GP, which is currently being evaluated in a phase I clinical trial (NCT05155332). We tested whether the dynamics of viral infection and therapeutic response could be directly assessed within our multicellular 3D melanoma-on-chip (Figure 5a). In conventional 2D cultures, MUG-Mel2 cells infected with VSV-GP-ZsGreen at different multiplicities of infection (MOI) exhibited efficient, homogeneous viral infection and reporter gene expression, evidenced by increasing ZsGreen fluorescence after 24 h (Figure 5b). Flow cytometric analysis confirmed productive viral replication under all MOIs (Supplementary Figure S6). Additionally, substantial oncolysis was observed with more than 80% loss of tumor cell viability in an MOI-dependent manner after 48 h (Figure 5c). While these findings demonstrate high intrinsic susceptibility of MUG-Mel2 cells to VSV-GP, conventional monolayer cultures fail to capture other factors known to influence the efficacy of oncolytic virotherapy in vivo, thereby overestimating the oncolytic effect in vivo. Thus, in the next step, we evaluated viral responses in the 3D bioprinted melanoma-on-chip models, which allowed us to assess the influence of the microenvironment on interactions between virus and tumor cells. In contrast to the rapid, homogeneous infection observed in 2D culture, ZsGreen fluorescence was not detectable in melanoma tumors after 24 h, but became apparent after 48 h, indicating delayed viral infection and propagation within the reconstructed 3D tissue environment (Figure 5d, Supplementary Figure S6a). Confocal fluorescence microscopy of infected melanoma-on-chip demonstrated viral infection almost exclusively in melanoma tumors (shown in red), while keratinocytes and fibroblasts in the surrounding skin tissue were not affected. Viral spread within the tumor was heterogeneous and did not uniformly cover the entire tumor spheroid (Figure 5e). Brightfield and fluorescence imaging further revealed progressive disruption of tumor architecture during treatment, accompanied by a reduction in spheroid size. Quantitative image analysis showed continuous growth of untreated spheroids, whereas VSV-GP treatment significantly decreased spheroid area (Figure 5f). Responses nevertheless varied among individual spheroids: while some underwent efficient lysis following infection, others displayed only transient ZsGreen expression. Overall a significant difference was observed when compared to untreated mock after 72 hours post infection (Figure 5g). To investigate the surrounding tissue, mean fluorescence intensity was quantified by fluorescence microscopy at 0, 24, 48, and 72 h in the region of interest (ROI) encompassing the MUG-Mel2 spheroid and in the surrounding non-tumor-bearing tissue. Comparison of mock-treated and VSV-GP-ZsGreen-treated chips revealed no increase in ZsGreen fluorescence intensity in the surrounding tissue, indicating an absence of detectable productive viral infection in fibroblasts and keratinocytes. (Figure 5h). These findings demonstrate that VSV-GP efficiently infects melanoma cells in both conventional 2D cultures and the engineered melanoma-on-chip tissue model. However, melanoma tumors embedded into a complex skin-TME exhibited delayed viral infection and tumor cell-restricted viral spread, and heterogeneous therapeutic responses, resulting in partial rather than complete tumor destruction. Taken together, these findings demonstrate that the tissue microenvironment can substantially influence viral infection, spread, and therapeutic efficacy, highlighting the importance of physiologically relevant tumor models for evaluating therapeutic responses.

**Figure 5.**
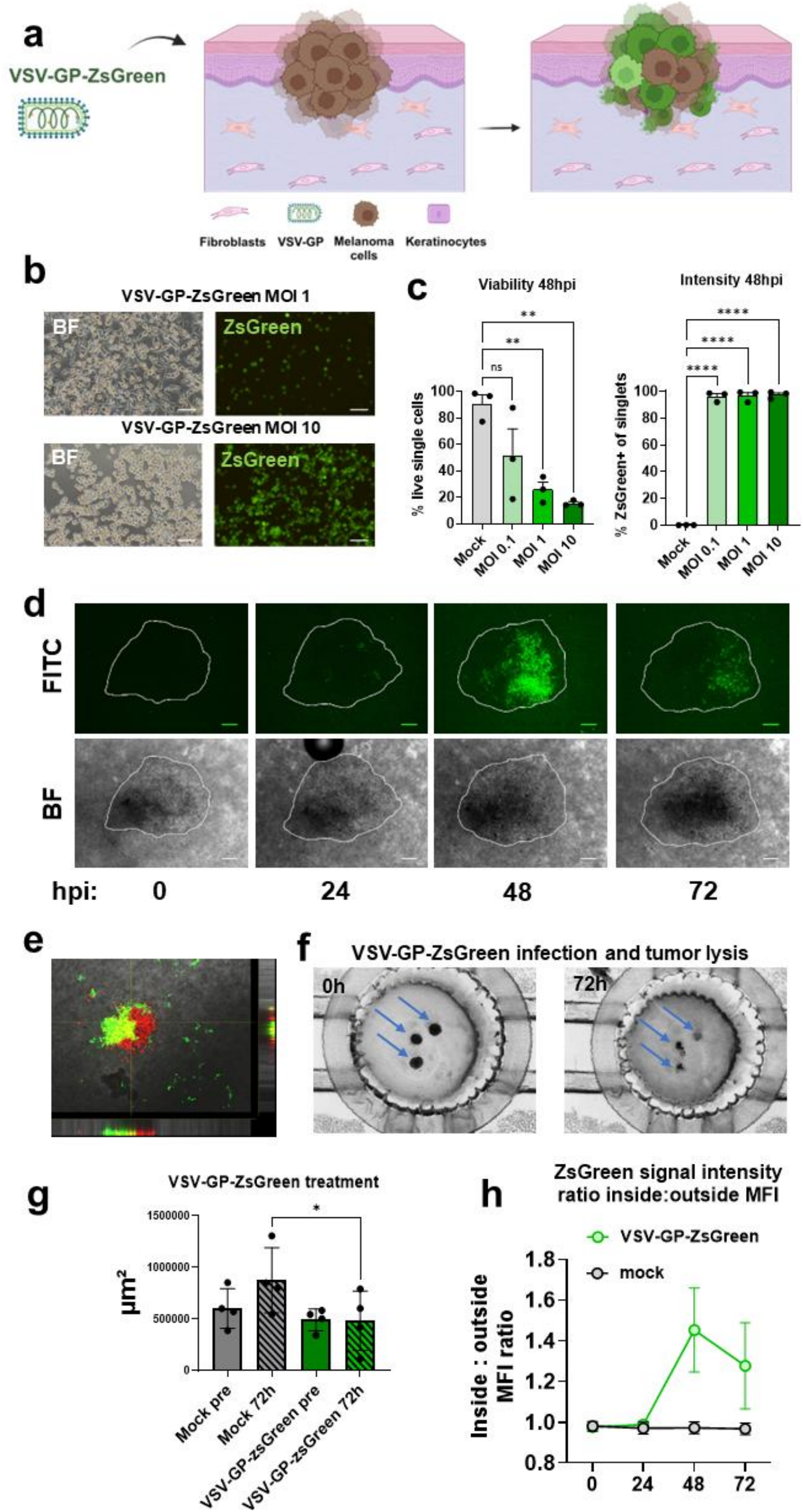
Melanoma-on-chip resembles a complex TME and is a suitable platform for oncolytic virus testing. (a) Schematic of the melanoma-on-chip model, illustrating the dermal compartment, the MUG-Mel2 melanoma spheroid, keratinocyte layer, and VSV-GP-ZsGreen virus. (b) Representative brightfield (BF) and GFP fluorescence microscopy images of MUG-Mel2 cells (2D culture) infected with VSV-GP-ZsGreen at MOI 1 and MOI 10 after 24 h. (c) Flow cytometric quantification after 48 h VSV-GP-ZsGreen infection at increasing multiplicities of infection (MOI; mock, 0.1, 1, and 10). Percentage of living single cells and ZsGreen-positive cells. Data are presented as mean ± SD (n = 3). Statistical analysis was performed using one-way ANOVA followed by Dunnett’s multiple comparisons test relative to the mock control (**P = 0.003; 0.076; ****P < 0.0001). (d) Representative fluorescence (top) and brightfield (bottom) images of melanoma-on-chip model at 0, 24, 48, and 72 h after VSV-GP-ZsGreen treatment, with the melanoma spheroid outlined, showing heterogeneous but progressive spread of ZsGreen signal within the tumor region. (e) Representative confocal image of MUG-Mel2 turboRFP melanoma cells 48 h post-infection (hpi) with VSV-GP-ZsGreen, MOI 10. (f) Macroscopic images of melanoma-on-chip model before and after VSV-GP-ZsGreen treatment (72 h). (g) Quantification of spheroid area (µm²) in melanoma-on-chip model comparing mock and VSV-GP-ZsGreen-treated conditions (*P = 0.0103). (h) Quantification of ZsGreen mean fluorescence intensity (MFI) over time (0, 24, 48, 72 h) inside versus outside the MUG-Mel2 spheroid region, comparing mock and VSV-GP-ZsGreen-treated chips; n=3 biological replicates from one representative experiment.

## Discussion

We developed a 3D bioprinted melanoma-on-chip platform that integrates a full-thickness human skin with spatially positioned melanoma spheroids right at the epidermal-dermal junction in a microfluidic chip to study the effect of novel melanoma therapeutics without animal experimentation. To generate the skin equivalents in customized laser-fabricated microfluidic chips, we combined extrusion printing for channel structures with multi-printhead and microjet printing for channel, dermal and epidermal layers in a single printing procedure. This enabled a medium-throughput production with short tissue printing times.

Tuning the optimal dermis stiffness proved fundamental for cell morphology and differentiation. GelMA concentrations were compared and showed that this significantly affected dermal fibroblast organization and morphology.

The stiffer formulation reduced fibroblast stretching but induced proper epidermis differentiation. Softer matrices resulted in a thinner and less organized epidermis, while the fibroblasts maintained an elongated, spread morphology indicative of healthy cell morphology. This is consistent with literature showing that dermal stiffness strongly influences epidermal mechanical properties and shape [27]. However, with 7.5% GelMA and 10% collagen for dermis composition, we observed proliferating keratinocytes above the basal layer in the epidermis. We concluded that the differentiation conditions were still not optimal. Therefore, we added an additional daily drying step, which ultimately produced a skin-on-chip model with a fully stratified epidermis that strongly expressed the differentiation markers CK1/10 and involucrin and predominantly Ki67 staining in the basal layer of the epidermis.

Successful barrier function of the 3D bioprinted skin was demonstrated by a lucifer-yellow permeability assay. To visualize different cell types and their distribution in the living 3D model, we transduced them with retroviruses coding for different fluorescent proteins and imaged them by multi-color confocal fluorescence microscopy. Fibroblasts and keratinocytes remained highly viable and fluorescent throughout the differentiation time, and embedded melanoma cells, despite their strong pigmentation, were detectable. Therefore, our model can also be used in the future to directly assess the impact (and unwanted potential side effects) of systemically applied melanoma therapeutics, as it allows assessment of skin cell dynamics during differentiation and treatment. Thus, we developed a membrane-free, full-thickness skin-on-chip model that replicates skin structure and key elements, including a highly differentiated epidermal layer.

Over the past years, numerous advanced melanoma tissue models have emerged (reviewed in [12]), ranging from organotypic skin equivalents to bioprinted constructs and microfluidic platforms. These efforts clearly reflect a growing recognition that more predictive human cell-based models beyond conventional cell culture systems and animal testing are needed. However, most existing systems capture only selected aspects of the disease. The challenge has been to develop a platform that simultaneously integrates complete epidermal maturation, precise spatial positioning of the tumor, robust long-term culture stability, and therapeutic evaluation capabilities within a single construct and with short fabrication timelines.

Recently Vázquez-Aristizabal and colleagues developed a medium-throughput model for melanoma in the OrganoPlate [13]. They developed a three-cell-type system using a high-capacity microfluidic device, placing melanoma tumor spheres inside a hydrogel derived from decellularized pig dermis, positioned next to channels lined with human endothelial and lymphatic cells. This approach highlights the value of tissue-derived matrices and demonstrates the feasibility of vascularized tumor chip testing at reasonable throughput. However, the model is confined to the dermal compartment and lacks an epidermal layer, which restricts its ability to recapitulate the full-thickness skin architecture, crucially, the tumor niche located at the junction between dermis and epidermis, where melanomas originate. Additionally, the constructs survive only for ten days and two key stromal components, the lymphatic endothelium and the decellularized scaffold itself are derived from murine and porcine sources. Yang et al. [14] demonstrated that patient-derived organoids with preserved immune components can respond to established BRAF-targeted therapies. However, their system lacks epidermal architecture, and the melanoma is not located at the epidermal-dermal junction where it naturally arises. Barros et al. [15] showed that a bioprinted melanoma model can support microneedle-based therapy delivery. Their work focused on transdermal drug penetration rather than mimicking the TME dynamics.

We addressed these gaps by developing a human 3D bioprinted skin model comprising a fibroblast-populated dermis that mimics human dermal thickness and stiffness and a multilayer differentiated epidermis. Critically, we positioned melanoma spheroids directly at the epidermal-dermal boundary, where melanomas arise *in vivo*. Our printing procedure combined different bioprinting techniques and produces twelve melanoma models in one run with printing times of about two minutes per chip. The tissues can be cultured for up to three weeks with efficient epidermis differentiation. Because the platform reconstructs the TME at realistic tissue scale and thickness rather than within miniaturized chip channels, it could enable detailed spatial and histological analysis of tumor-stroma interactions. This tissue-scale engineering also allows us to observe phenomena that miniaturized systems cannot capture. For detailed studies, we selected an NRAS-mutated melanoma cell line which was established recently rather than the more studied BRAF-mutated line used by other groups [28]. NRAS melanoma has fewer treatment options and more aggressive tumor growth, making it an interesting target for experimental therapeutic approaches.

Understanding how tissue architecture influences therapeutic outcomes is crucial for developing novel anti-cancer treatment strategies. In standard monolayer culture, MUG-Mel2 melanoma cells exposed to the oncolytic virus VSV-GP-ZsGreen showed rapid homogeneous infection with approximately 80% cell death within 48 h. The virus appeared effective at eliminating melanoma cells, since there was no heterogeneity in the tumor cell population. However, inside our 3D tissue model, the situation changed. The multicellular TME of the melanoma-on-chip model was reflected in slower virus dissemination, apparent by a delayed onset of ZsGreen signal at around 48 h post infection. In a perfused full-thickness skin-on-chip, treatment responses likewise developed with a delay relative to 2D monocultures, consistent with the more physiological response kinetics observed here [29]. Further, virus spread was limited to tumor cells, yet proceeded not uniformly like in the 2D system. Similar findings have been reported in a perfused 3D microphysiological system, in which oncolytic VSV-GFP replication was predominantly detected in tumor cells, while stromal fibroblasts and endothelial cells showed no detectable viral replication. These observations highlight the influence of the surrounding tissue microenvironment on the spatial restriction of viral infection [30]. The lytic efficacy of VSV-GP-ZsGreen (shrinkage or destruction of tumor spheroids) showed marked heterogeneity in the 3D model. These observations align with Kimpel et al. [21] who showed that VSV-GP efficiently lysed all tested melanoma cell lines in 2D culture, yet exhibited significantly reduced efficacy in mouse models, with only a few animals achieving long-term tumor remission. The authors attributed this discrepancy to intact type I interferon signaling in tumor cells, which effectively restricted viral replication in vivo. Our 3D bioprinted melanoma-on-chip model provides a microenvironment that more closely resembles the in vivo situation, incorporating physiologically relevant extracellular matrix components and cell-cell interactions. By integrating fibroblasts and keratinocytes within the 3D construct, we better recapitulate the complex TME that governs oncolytic viral kinetics. Therefore, our melanoma-on-chip could aid in developing a more potent in vivo treatment strategy against melanoma in the future.

The presented disease platform has limitations that we are also currently addressing in ongoing projects: Our platform does not include immune cell populations, which play an important role in anti-tumor immunity and especially oncolytic virotherapy. The incorporation of exogenously supplied effector T cells into organotypic melanoma-in-skin models has previously been demonstrated through intradermal cell injection [31]. However, the establishment of an immune compartment within the tissue construct, allowing immune cells to be introduced through a perfusable vascular network, remains an important step towards greater physiological complexity. In our platform, this approach is currently being investigated to enable immune cell recruitment and interaction with the tumor microenvironment via the vascular channels. Further, we used immortalized cells for this study to achieve sufficient cell numbers for printing multiple tissue equivalents rather than primary cells or tumor material, thereby missing the full genetic diversity of actual patient-derived specimens. Despite these limitations, the current platform clearly demonstrates that 3D tissue architecture fundamentally alters therapeutic responses in ways that correlate with animal models, supporting its potential as a physiologically relevant system for preclinical evaluation and as a meaningful step toward replacing animal experiments.

## Conclusion

The presented 3D bioprinted melanoma-on-chip approach combines single-step fabrication of a channel-containing skin equivalent with custom-designed, laser-manufactured fluidic chips, enabling long-term culture of the melanoma-on-chip model. Precise spatial localization of melanoma spheroids at the epidermal-dermal junction sets this melanoma model apart from currently existing systems. The successful application of oncolytic virus treatment demonstrates the feasibility of investigating therapeutic responses within a complex, 3D skin microenvironment. Unlike conventional subcutaneous melanoma animal models, this platform incorporates key structural features of human skin and offers a complementary approach to studying tumor– tissue interactions during treatment. Furthermore, its applicability to patient-derived melanoma cells, as demonstrated using MUG-Mel2 cells, highlights its potential for evaluating therapeutic responses in a patient-specific tissue context and supporting future precision medicine approaches.

## Materials and Methods

Unless otherwise stated, chemicals and reagents were obtained from Merck.

### CELL LINES AND CULTURE CONDITIONS

Human dermal fibroblast cells (HDFs; ATCC®, PCS-201-010), human keratinocytes (HaCaT; Cytion GmbH, Heidelberg, Germany) were cultured in DMEM with high glucose (Thermo Fisher Scientific, Waltham, USA). HUVEC cells (ATCC® CRL-1730™) were cultured with Ham’s F-12 K (Kaighn’s) medium (Thermo Fisher Scientific, Waltham, USA) supplemented with 30 µg/ml endothelial cell growth supplement (ECGS), and 100 µg/ml heparin.

BHK-21 cells (Elabscience #EP-CL-0034) were maintained in Glasgow minimum essential medium-BHK-21 (G-MEM BHK21 #2458821) (Gibco, Paisley, Scotland, UK) and 5% tryptose phosphate broth (Gibco, Paisley, Scotland, UK). MUG-Mel2 and MUG-Mel3 patient-derived melanoma cells (Medical University Graz, Austria) and A375 melanoma cells (ATCC®, CRL-1619) were maintained in RPMI 1640.

All media contained 10% fetal bovine serum (Gibco, Paisley, Scotland, UK), 100 U/mL penicillin, 100 µg/mL streptomycin, and 2 mM L-glutamine (Lonza, Basel, Switzerland). Cells were cultured at 37 °C in a humidified atmosphere with 5% CO2 and detached at 80% confluence using Accutase (Innovative Cell Technologies). MUG-Mel2 and MUG-Mel3 cell isolation was approved by the local ethics committee at Medical University of Graz (vote #18 192ex06/07). Routine mycoplasma testing was performed using the Venor®GeM Advance Mycoplasma Detection Kit (Minerva Biolabs GmbH, Berlin, Germany) according to the manufacturer’s instructions. The assay is based on conventional endpoint PCR, and amplified PCR products were analyzed by agarose gel electrophoresis.

### MICROFLUIDIC CHIP FABRICATION

Fabrication of fluidic chips was described previously [16] briefly, microfluidic devices were designed in CorelDraw Graphics Suite 2020 (Corel Corporation, Ottawa, Ontario, Canada) and laser cut from 3 mm PMMA sheets using a Mini Epilog Laser system (Epilog Laser, Colorado, USA). Chips were bonded to glass microscopy coverslips (21 × 26 mm, thickness 0.16 mm; Menzel-Gläser) using biocompatible double-sided adhesive tape (ARcare 90106NB, Adhesive Research Inc., Pennsylvania, USA). Chips were washed in an ultrasonic bath for 15 min, boiled twice in distilled water for 5 min, and sterilized overnight in 70% ethanol. After drying, chips were exposed to UV-C light for 20 min and plasma-treated (Atto Plasma System, Diener Electronic) for 10 min before bioprinting. For stereolithography (SLA) resin chips, a consumer DLP printer (Anycubic Photon D2) was used with biocompatible resin (Bio-Med Clear, Liqcreate, Netherlands). Printed chips were washed in isopropanol, post-cured under 405 nm violet light for 30 min and sterilized following the same protocol as PMMA chips.

### GELMA SYNTHESIS

GelMA was synthesized following the protocol of [32] with modifications. Briefly 60 g gelatin was dissolved in 600 mL of 0.25 ᴍ carbonate-bicarbonate buffer (10% w/v) at 50 °C with stirring at 600 rpm. Methacrylic anhydride was added at a ratio of 6 g per 60 g gelatin and the solution was stirred for 90 min at 50 °C. The solution was transferred to dialysis tubing (6-8 kDa MWCO) and dialyzed against deionized water for 4 days at 50 °C. The GelMA solution was lyophilized and stored at -80 °C.

### MECHANICAL CHARACTERIZATION OF GELMA HYDROGELS

GelMA hydrogels 5%, 7.5% or 10% w/v, were each supplemented with 10% w/v collagen (Arthro Kinetics Biotechnology, Krems an der Donau, Austria) with a degree of functionalization of approximately 90%, as determined by the TNBS assay. The bioink was characterized using an ElastoSens Bio rheometer (Rheolution Inc., Montréal, QC, Canada) at 20 °C. Measurements were performed every 10 s using: 5 min without 405 nm LED light (0% intensity), 10 min with 405 nm LED light (70% intensity), and 5 min without light.

### 3D BIOPRINTING

GelMA/LAP bioink was prepared in DMEM (5% or 7.5% GelMA, 0.25% LAP) by incubation at 37 °C for 3 h and stored at 4 °C overnight. Thermo-reversible Pluronic F127 (35% in 0.9% NaCl) was first printed as sacrificial ink at 160 kPa to generate hollow channels. HDFs were mixed 1:1 (v/v) with prewarmed GelMA/LAP bioink to achieve 8.75 × 10^6^ cells/mL. Collagen I (Arthro Kinetics Biotechnology, Krems an der Donau, Austria) (10% w/v) was added immediately before printing. HDF-laden hydrogels were printed at 37 °C by microvalve jetting at 12 kPa in three sequential layers. Each layer was polymerized by photo-crosslinking (10 s at 365 nm, 360 mW) using the integrated LED. MUG-Mel2 melanoma spheroids (1– 5 per chip) were manually transferred onto the top dermis layer before final crosslinking. For epidermal constructs, HaCaT keratinocytes (12 × 10^6^ cells/mL) were suspended in PVP/DMEM (2.5% PVP) and deposited in two layers at 30 kPa. Constructs were crosslinked for 30 sec at 405 nm LED light. After 24 h in submerged conditions, constructs were raised to air-liquid interface (ALI) and cultured for 2–3 weeks in KGM (3:1 DMEM:F12) supplemented with 10 ng/mL rhEGF (ImmunoTools GmbH, Friesoythe, Germany), 0.4 µg/mL hydrocortisone, 50 µg/mL L-ascorbic acid, 24 µg/mL adenine, 1× ITS (Gibco, Paisley, Scotland, UK). Daily controlled air-drying (30 min in biosafety hood) promoted epidermal differentiation.

### MELANOMA SPHEROID FORMATION

MUG-Mel2, MUG-Mel3 and A375 spheroids were generated by seeding 10,000 cells per well into 96-well V-bottom plates (Corning, NY, USA) followed by centrifugation at 300 × *g* for 5 min. Cells were cultured for 3 days to allow spheroid formation. Spheroid diameter (∼400 µm) was verified by brightfield microscopy (Axiovert 200M, Zeiss).

### VECTORS, RETROVIRAL AND LENTIVIRAL INFECTIONS

The retroviral vector pLIB-mCherry-iresPuro was used to infect HaCaT keratinocytes (HaCaT/mCherry) and pQXCIN TurboRFP-Puro was used for transducing MUG-Mel2. Retroviral vectors pLIB-hTert-CMVpuro and pQCXIN-EYFP-iresNeo were transfected into Phoenix packaging cells using Lipofectamine 2000 (Thermo Fisher Scientific, Waltham, MA, USA) and virus supernatant was used to immortalize HDFs and HUVEC cells and label with EYFP (HDF/hTert-EYFP, HUVEC/hTert-EYFP) as described previously [16, 33]. The lentiviral vector pHR SFFV emiRFP670 ires-puro was constructed in the following way: The coding sequence of emiRFP670 was PCR-amplified from pemiRFP670-N1 (Addgene #372972, [25]), the iresPuro sequence was amplified from pLIB-MCS-iresPuro [34], assembled using Gibson-Cloning (New England Biolabs, Ipswitch, MA, USA) and inserted into the lentiviral vector pHR SFFV eGFP which had been digested with BamHI and NotI to remove the eGFP coding sequence.

### HUMAN SKIN SAMPLES

Healthy human skin was obtained as residual tissue from plastic surgery at Medical University of Innsbruck. Samples were fixed in 4% formaldehyde and paraffin-embedded and used as reference tissue for histological and immunohistochemical comparison. Tissue collection was approved by the Ethics Committee of the Medical University of Innsbruck AN5003 and all donors provided written informed consent.

### HISTOLOGY AND IMMUNOHISTOCHEMISTRY

Paraffin-embedded samples were sectioned and stained with hematoxylin and eosin (H&E) for histological evaluation. Immunohistochemical staining was performed using antibodies against Ki67, AE1-AE3 (pan-cytokeratin), S100, and vimentin, and staining procedures were performed according to the manufacturer’s instructions (Ventana Roche).

For immunofluorescence, sections were deparaffinized in ROTI®Histol (Carl Roth GmbH + Co., Karlsruhe, Germany) (2 × 5 min), rehydrated through a descending ethanol series, and treated with 3% H₂O₂ in PBS for 10 min. Antigen retrieval was performed in 10 mM citrate buffer using a microwave (20 min), followed by blocking for 60 min at room temperature. Sections were incubated overnight at 4 °C with primary antibodies against cytokeratin 1/10 (CBL266; 1:50) and involucrin (Abcam, Cambridge, UK; ab53112; 1:75).

### PROTEIN EXTRACTION

Tissue samples were lysed in 320 µL Skin Lysis Buffer per sample supplemented with Pierce™ Protease and Phosphatase Inhibitor Mini Tablets (Thermo Fisher Scientific) at a final 1× concentration. Samples were placed on ice, homogenized using an Ultra-Turrax (IKA, Staufen, Germany) until completely dissolved and then sonicated twice for 9 s at 70% amplitude using a Sonoplus HD 2070 apparatus (Bandelin Electronic, Ebhausen, Germany). Lysates were incubated on a rotary wheel at 4 °C for 45 min and centrifuged at 18620 × g for 30 min at 4 °C. Supernatants (300 µL) were collected and mixed with 100 µL of 4× SSD loading dye 4× SSB sample buffer was prepared containing 8% (w/v) SDS, 250 mM Tris-HCl (pH 6.8), 40% (v/v) glycerol (Carl Roth GmbH + Co., Karlsruhe, Germany), and 1 mg/mL bromophenol blue. 5% β-mercaptoethanol (Bio-Rad Laboratories, Hercules, California, USA) was freshly added and samples were stored at -20 °C.

### TISSUE LYSIS

A lysis buffer was prepared to a final volume of 200 mL containing 50 mM HEPES/NaOH (pH 7.5), 150 mM NaCl, 5 mM EDTA, 10% (v/v) glycerol, 1 mM DTT, 1% (v/v) Triton X-100, 1% (w/v) SDS, and 1% (v/v) IGEPAL.

### IMMUNOBLOTTING

Cytokeratin 1/10 protein levels were analyzed using an automated capillary electrophoresis system (Jess Simple Western^TM^, R&D systems, Bio-Techne, Minneapolis, MN, USA). Lysates were loaded onto 12–230 kDa separation capillary modules (Bio-Techne, Cat. No. SM-W001). Capillaries were probed with primary antibodies against cytokeratin 1/10 at 1:25 dilution, followed by HRP-conjugated goat anti-mouse secondary antibody (Bio-Techne, Cat. No. 042-205). Vinculin (loading control) was detected with anti-vinculin antibody (Cell Signaling Technology, Cat. No. 13901S, clone E1E9V) at 1:50000 dilution, followed by HRP-conjugated goat anti-rabbit secondary antibody (Bio-Techne, Cat. No. 042-206). Signal quantification was done in Compass for Simple Western software (v7.1.0, Bio-Techne).

Skin Barrier Function Assay: Barrier function assay was performed following the protocol of [35] with slight adaptations, constructs were treated with or without 1% sodium dodecyl sulfate (SDS) for 2 h prior to assay (positive control). Lucifer Yellow CH dilithium salt (1 mg/mL in PBS) was applied topically (10 µL) onto the epidermis at ALI on day 14. Lucifer Yellow fluorescence increase in culture medium was measured after 1 h and 2 h. For quantification, 10 µL culture medium was diluted 1:5 in FluoroBright DMEM (Thermo Fisher Scientific) and transferred to black 96-well plates (Corning, NY, USA). Fluorescence was measured (Ex 428 nm, Em 536 nm) using a Tecan Spark plate reader (Tecan Trading AG, Switzerland).

### FLOW CYTOMETRY

MUG-Mel2 cells were seeded in 24-well plates and treated with three different viral dilutions for 48 h. Cells were analyzed for ZsGreen reporter expression and viability using 1:1000 ToPro3 dye (Invitrogen, Cat. No. R37170) on a MACSQuant 16 Flow Analyzer (Miltenyi Biotec B.V. & Co. KG, Bergisch Gladbach, Germany). After gating for singlet events, the percentage of ZsGreen-positive events or ToPro3-negative (live) events was determined from technical triplicates in three independent experiments. Data were analyzed using FlowJo software (v10, BD Biosciences) and plotted using GraphPad Prism (v10).

### MELANOMA-ON-CHIP INFECTION

Melanoma-on-chip constructs were infected with VSV-GP-ZsGreen virus at MOI 10 (calculated on ∼60,000 MUG-Mel2 cells per construct, 6 × 10^5^ TCID50 well^-1^). 3D bioprinted melanoma-on-chips were dried for 10 min, then infected by adding 100 µL complete growth medium containing VSV-GP-ZsGreen. Plates were transferred to the incubator undisturbed for 1 h, before 1 mL complete medium was added. Constructs underwent controlled daily drying (30 min in biosafety hood) and were monitored by fluorescence microscopy.

### IMAGE QUANTIFICATION OF INFECTED MELANOMA-ON-CHIP

Fluorescence images were acquired on an Olympus IX70 microscope (Olympus) with 4× objective lens using identical exposure times. Raw images were processed in ImageJ (Fiji distribution, version 1.54g). For mean fluorescence intensity (MFI) quantification, regions of interest (ROIs) were manually drawn on 8-bit, flattened phase-contrast images to define inside (spheroid) and outside (background) areas. These ROIs were applied to raw FITC-channel images (8-bit, flattened), and mean gray values were extracted. MFI was plotted using GraphPad Prism (v11). Three melanoma-on-chip *(n = 3)* constructs from one representative experiment were analyzed at four time points per condition. Melanoma growth was assessed at day 1 and day 21 using a digital microscope (Keyence VHX-6000) at 20× magnification with a 4K image sensor. The projected area of individual melanoma spheroids was quantified in mm² at each time point. Three spheroids per chip were analyzed across seven independent experiments *(n = 7)*.

### VIRUS STOCK PRODUCTION AND TCID50 TITRATION

VSV-GP-ZsGreen virus was produced as previously described by [24] via a reverse genetics approach in a helper virus-free protocol (Witko et al 2006, PMID 16569439). Briefly, a ZsGreen cargo cassette was inserted into a parent DNA plasmid encoding all VSV-GP genes (N, P, M, LCMV-GP, L) on the 5^th^ position between LCMV-GP and L, analogous to the generation of VSV-GP-GFP described in [24] via restriction enzyme-mediated digestion and re-ligation via gibson assembly (NEBuilder HiFi DNA Assembly kit, NEB #E2623). Functional VSV-GP-ZsGreen virus was recovered via transfection of VSV-GP-ZsGreen plasmid into HEK293T cells together with helper plasmids.

Infectious virus was propagated on hamster BHK-21 cells and plaque-purified twice before concentration, aliquots were stored on -80 °C. Titers were assessed as TCID50/mL on BHK-21 cells. In short, half-log serial dilutions of virus stock aliquots were prepared in complete BHK-21 medium. 100 µL were then added in 8 technical replicates onto wells containing a confluent BHK-21 cell culture and incubated for five days, until a cytopathic effect was visible. TCID50/mL values were calculated according to the Spearman-Kärber method. Fresh aliquots were used for each experiment.

### LIVE CELL FLUORESCENCE MICROSCOPY AND CONFOCAL IMAGING

Immunofluorescence staining, fibroblast distribution, and melanoma were imaged in an Axiovert 200M microscope (Zeiss, Oberkochen, Germany). Melanoma spheroid area and diameter were quantified from brightfield and fluorescence images using ImageJ (Fiji distribution) by manual outlining of spheroid boundary in 3–5 spheroids per condition and time point. Fibroblast circularity was determined from EYFP-expressing HDF images using an ImageJ macro. Individual cells were identified using the Frangi filter (parameters: scale = 10, spacing = 1), thresholding and the ‘Analyze particles’ tool, excluding debris below 50 µm^2^ and large aggregates above 1500 µm^2^. Circularity was identified for each ROI using the shape descriptor function and averaged across each image. The final analysis contains at least 6 images per condition and time point from 3 independent experiments. Confocal 3D stacks of skin tissue were acquired at an FV4000 confocal microscope (Evident, Japan) equipped with 10 lasers for multi-color excitation/imaging also in the near infrared spectrum. Overview 3D images acquired with a 4× objective were stitched and visualized in Fluoview^TM^ software (Evident, Japan).

### STATISTICAL ANALYSIS

GraphPad Prism software (v11; GraphPad Software, San Diego, California, USA) was used for all statistical analyses. Comparisons between two groups were performed using unpaired Student’s t-tests or Welch’s t-tests (for unequal variances). Comparisons among more than two groups used one-way analysis of variance (ANOVA) with Dunnett’s multiple comparisons test relative to mock controls, or two-way ANOVA for factorial designs. Differences were considered statistically significant at *P < 0.05*. Results are reported as mean ± standard deviation (n ≥ 3 unless otherwise specified). Significance levels are denoted as: ns (not significant), *\*P < 0.05, **P < 0.01, ***P < 0.001, ****P < 0.0001*.

Supplementary Materials: The Supplementary Materials are available online at:

## Acknowledgments

We thank B Moser, M Binder, M. Abenthung, N. Kühner, K. Heinz, V. Jeller, V. Hackl, D. Nothdurfter and N. Haidenhofer for their technical support. We further thank Colin McGuckin and Nico Forraz (CTI-Biotech) and the European Network for Skin Engineering and Modeling (CA21108) for valuable discussions. Funding was received from the Dr Johannes und Herta Tuba-Stiftung, Austrian Science Fund Special Research Group FG15 (10.55776/FG15), Austrian Science Fund Project PAT1161725 (10.55776/PAT1161725), the Austrian Research Promotion Agency (Bridge-1 880666), the intramural PhD funding program of the Medical University Innsbruck CONNECT, and the Tirol-Kliniken GmbH.

## Author Contributions

Study design and concept: J H, M J A, P S, A K Data acquisition and analysis: A K, N K, E B, A A, E H, G W, B R, D O T, P S, J H, M J A Drafting the manuscript: A K, J H, M J A

## Table of Contents

A full-thickness human skin is bioprinted directly inside a microfluidic chip, with melanoma spheroids positioned at the epidermal-dermal junction. Tuning the dermal matrix stiffness controls epidermal maturation and barrier formation and yields tumor-bearing tissues that remain stable for 21 days. Oncolytic virus infection spreads slowly and stays confined to tumor cells, revealing how tissue architecture shapes therapeutic response.

## References

1. M. Arnold, D. Singh, M. Laversanne, et al., "Global Burden of Cutaneous Melanoma in 2020 and Projections to 2040," JAMA Dermatology 158, no. 5 (2022): 495–503, 10.1001/jamadermatol.2022.0160.

2. E. Hodis, I. R. Watson, G. V. Kryukov, et al., "A Landscape of Driver Mutations in Melanoma," Cell 150, no. 2 (2012): 251–263, 10.1016/j.cell.2012.06.024.

3. J. Larkin, V. Chiarion-Sileni, R. Gonzalez, et al., "Five-Year Survival with Combined Nivolumab and Ipilimumab in Advanced Melanoma," New England Journal of Medicine 381, no. 16 (2019): 1535–1546, 10.1056/NEJMoa1910836.

4. D. B. Johnson, A. M. Menzies, L. Zimmer, et al., "Acquired BRAF Inhibitor Resistance: A Multicenter Meta-Analysis of the Spectrum and Frequencies, Clinical Behaviour, and Phenotypic Associations of Resistance Mechanisms," European Journal of Cancer 51, no. 18 (2015): 2792–2799, 10.1016/j.ejca.2015.08.022.

5. T. N. Gide, J. S. Wilmott, R. A. Scolyer, and G. V. Long, "Primary and Acquired Resistance to Immune Checkpoint Inhibitors in Metastatic Melanoma," Clinical Cancer Research 24, no. 6 (2018): 1260–1270, 10.1158/1078-0432.CCR-17-2267.

6. S. Y. Lim, E. Shklovskaya, J. H. Lee, et al., "The Molecular and Functional Landscape of Resistance to Immune Checkpoint Blockade in Melanoma," Nature Communications 14, no. 1 (2023): 1516, 10.1038/s41467-023-36979-y.

7. H. Sikorski, M. A. Zmijewski, and A. Piotrowska, "Tumor Microenvironment in Melanoma: Characteristic and Clinical Implications," International Journal of Molecular Sciences 26, no. 14 (2025): 6778, 10.3390/ijms26146778.

8. J. Magrill, D. Moldoveanu, J. Gu, M. Lajoie, and I. R. Watson, "Mapping the Single Cell Spatial Immune Landscapes of the Melanoma Microenvironment," Clinical & Experimental Metastasis 41, no. 4 (2024): 301–312, 10.1007/s10585-023-10252-4.

9. S. Di Blasio, G. F. van Wigcheren, A. Becker, et al., "The Tumour Microenvironment Shapes Dendritic Cell Plasticity in a Human Organotypic Melanoma Culture," Nature Communications 11, no. 1 (2020): 2749, 10.1038/s41467-020-16583-0.

10. F. Bojin, A. Robu, M. I. Bejenariu, et al., "3D Bioprinting of Model Tissues That Mimic the Tumor Microenvironment," Micromachines 12, no. 5 (2021): 535, 10.3390/mi12050535.

11. N. D. Daugaard, R. Tholstrup, J. R. Tornby, et al., "Characterization of Human Melanoma Skin Cancer Models: A Step Towards Model-Based Melanoma Research," Acta Biomaterialia 191 (2025): 308–324, 10.1016/j.actbio.2024.11.018.

12. K. Xu, J. Wu, Z. Li, et al., "The Application of 3D Cell Culture for Melanoma In Vitro Models," Acta Biomaterialia 209 (2026): 148–166, 10.1016/j.actbio.2025.11.039.

13. P. Vázquez-Aristizabal, I. Arriola-Alvarez, M. Henriksen-Lacey, et al., "Decellularized Dermis ECM-Based Melanoma-on-a-Chip Model with Integrated Lymphatic and Vascular Networks for High-Throughput Drug Testing," ACS Applied Bio Materials 9, no. 6 (2026): 2971–2979, 10.1021/acsabm.5c02329.

14. F. Yang, B. Shi, J. Zhao, et al., "Development of a Human Melanoma Organoid-Containing Skin-on-a-Chip Model for In Vitro Drug Testing and Personalized Medicine," Biofabrication 18, no. 3 (2026), 10.1088/1758-5090/ae8e03.

15. N. R. Barros, R. Kang, J. Kim, et al., "A Human Skin-on-a-Chip Platform for Microneedling-Driven Skin Cancer Treatment," Materials Today Bio 30 (2025): 101399, 10.1016/j.mtbio.2024.101399.

16. D. Nothdurfter, C. Ploner, D. C. Coraça-Huber, et al., "3D Bioprinted, Vascularized Neuroblastoma Tumor Environment in Fluidic Chip Devices for Precision Medicine Drug Testing," Biofabrication 14, no. 3 (2022): 035002, 10.1088/1758-5090/ac5fb7.

17. J. Hagenbuchner, D. Nothdurfter, and M. J. Ausserlechner, "3D Bioprinting: Novel Approaches for Engineering Complex Human Tissue Equivalents and Drug Testing," Essays in Biochemistry 65, no. 3 (2021): 417–427, 10.1042/EBC20200153.

18. A. Muik, L. J. Stubbert, R. Z. Jahedi, et al., "Re-Engineering Vesicular Stomatitis Virus to Abrogate Neurotoxicity, Circumvent Humoral Immunity, and Enhance Oncolytic Potency," Cancer Research 74, no. 13 (2014): 3567–3578, 10.1158/0008-5472.CAN-13-3306.

19. C. Urbiola, F. R. Santer, M. Petersson, et al., "Oncolytic Activity of the Rhabdovirus VSV-GP against Prostate Cancer," International Journal of Cancer 143, no. 7 (2018): 1786–1796, 10.1002/ijc.31556.

20. L.-M. Schreiber, C. Urbiola, K. Das, et al., "The Lytic Activity of VSV-GP Treatment Dominates the Therapeutic Effects in a Syngeneic Model of Lung Cancer," British Journal of Cancer 121, no. 8 (2019): 647–658, 10.1038/s41416-019-0574-7.

21. J. Kimpel, C. Urbiola, I. Koske, et al., "The Oncolytic Virus VSV-GP Is Effective against Malignant Melanoma," Viruses 10, no. 3 (2018): 108, 10.3390/v10030108.

22. E. Hastie and V. Z. Grdzelishvili, "Vesicular Stomatitis Virus as a Flexible Platform for Oncolytic Virotherapy against Cancer," Journal of General Virology 93, no. 12 (2012): 2529–2545, 10.1099/vir.0.046672-0.

23. S. Rivera-Orellana, J. Bautista, D. Palacios-Zavala, et al., "Oncolytic Virotherapy and Tumor Microenvironment Modulation," Clinical and Experimental Medicine 25, no. 1 (2025): 256, 10.1007/s10238-025-01691-2.

24. C. Dold, C. Urbiola, G. Wollmann, et al., "Application of Interferon Modulators to Overcome Partial Resistance of Human Ovarian Cancers to VSV-GP Oncolytic Viral Therapy," Molecular Therapy – Oncolytics 3 (2016): 16021, 10.1038/mto.2016.21.

25. M. E. Matlashov, D. M. Shcherbakova, J. Alvelid, et al., "A Set of Monomeric Near-Infrared Fluorescent Proteins for Multicolor Imaging across Scales," Nature Communications 11, no. 1 (2020): 239, 10.1038/s41467-019-13897-6.

26. B. Rinner, G. Gandolfi, K. Meditz, et al., "MUG-Mel2, a Novel Highly Pigmented and Well Characterized NRAS Mutated Human Melanoma Cell Line," Scientific Reports 7, no. 1 (2017): 2098, 10.1038/s41598-017-02197-y.

27. E. Roig-Rosello, G. Dayan, S. Bovio, et al., "Dermal Stiffness Governs the Topography of the Epidermis and the Underlying Basement Membrane in Young and Old Human Skin," Aging Cell 23, no. 4 (2024): e14096, 10.1111/acel.14096.

28. U. Ben-David, B. Siranosian, G. Ha, et al., "Genetic and Transcriptional Evolution Alters Cancer Cell Line Drug Response," Nature 560, no. 7718 (2018): 325–330, 10.1038/s41586-018-0409-3.

29. S. Rhee, C. Xia, A. Chandra, et al., "Full-Thickness Perfused Skin-on-a-Chip with In Vivo-Like Drug Response for Drug and Cosmetics Testing," Bioengineering 11, no. 11 (2024): 1055, 10.3390/bioengineering11111055.

30. K. J. Lee, S. W. Lee, H. N. Woo, et al., "Real-Time Monitoring of Oncolytic VSV Properties in a Novel In Vitro Microphysiological System Containing 3D Multicellular Tumor Spheroids," PLoS ONE 15, no. 7 (2020): e0235356, 10.1371/journal.pone.0235356.

31. E. Michielon, L. A. King, T. Waaijman, et al., "An Organotypic Human Melanoma-in-Skin Model as an In Vitro Tool for Testing Vγ9Vδ2-T Cell-Based Immunotherapy," Immuno-Oncology and Technology 24 (2024): 100724, 10.1016/j.iotech.2024.100724.

32. H. Shirahama, B. H. Lee, L. P. Tan, and N.-J. Cho, "Precise Tuning of Facile One-Pot Gelatin Methacryloyl (GelMA) Synthesis," Scientific Reports 6, no. 1 (2016): 31036, 10.1038/srep31036.

33. J. Hagenbuchner, V. Obsilova, T. Kaserer, et al., "Modulating FOXO3 Transcriptional Activity by Small, DBD-Binding Molecules," eLife 8 (2019): e48876, 10.7554/eLife.48876.

34. M. J. Ausserlechner, C. Salvador, A. Deutschmann, et al., "Therapy-Resistant Acute Lymphoblastic Leukemia (ALL) Cells Inactivate FOXO3 to Escape Apoptosis Induction by TRAIL and Noxa," Oncotarget 4, no. 7 (2013): 995– 1007, 10.18632/oncotarget.953.

35. J. Holzknecht, S. Dubrac, S. Hedtrich, L. Galgóczy, and F. Marx, "Small, Cationic Antifungal Proteins from Filamentous Fungi Inhibit Candida albicans Growth in 3D Skin Infection Models," Microbiology Spectrum 10, no. 3 (2022): e0029922, 10.1128/spectrum.00299-22.

