## Supplemental Figures for "A Full-Thickness 3D Bioprinted Melanoma-on-Chip Platform for Preclinical Therapeutic Testing"

***Supplementary Figures:***

Supplementary Figure S1. Overview RegenHu 3D bioprinting setup and printing process 2

Supplementary Figure S2. Comparison of alternative microfluidic chip designs 3

Supplementary Figure S3. Effect of GelMA concentration on gel stiffness 4

Supplementary Figure S4. Vascularized skin-on-chip 5

Supplementary Figure S5. Comparison of different melanoma cell lines in 2D and 3D 6

Supplementary Figure S6. Representative VSV-GP-ZsGreen infected melanoma-on-chip 7

**a**

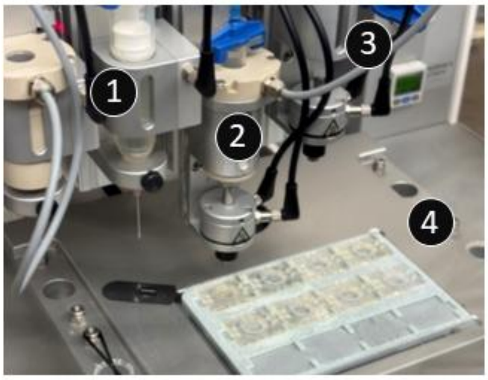


**b**

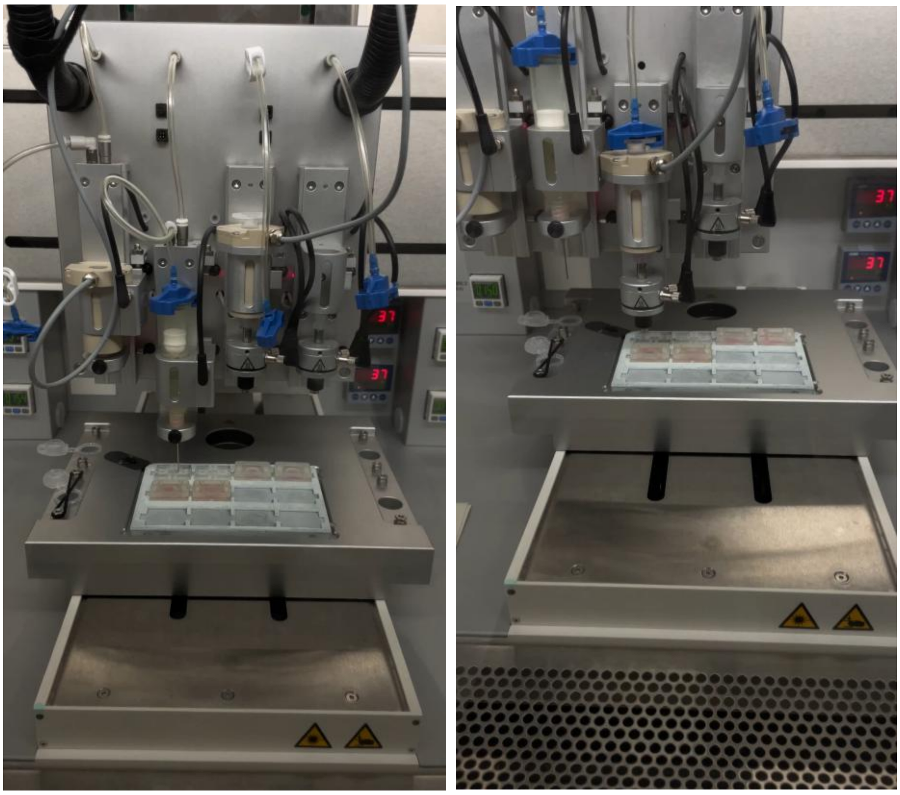

**Supplementary Figure S1. Overview RegenHu 3D bioprinting setup and printing process.** (a) Overview of the RegenHu bioprinting setup used for melanoma-on-chip fabrication. (1) Inkjet printhead for dispensing the Pluronic sacrificial channel. (2) Extrusion printhead for HDF (human dermal fibroblast) bioprinting, heated to 37°C. (3) Extrusion printhead for HaCaT (keratinocyte) bioprinting, heated to 37°C. (4) Platform holding the microfluidic chips during printing, accommodating up to 12 chips simultaneously. (b) Representative image of supplementary videos 1 and 2 of the bioprinting process, showing the printhead assembly depositing material onto the chip platform loaded with microfluidic chips; (left) Pluronic channel printing is shown; (right) 3D bioprinting of the dermis is shown.

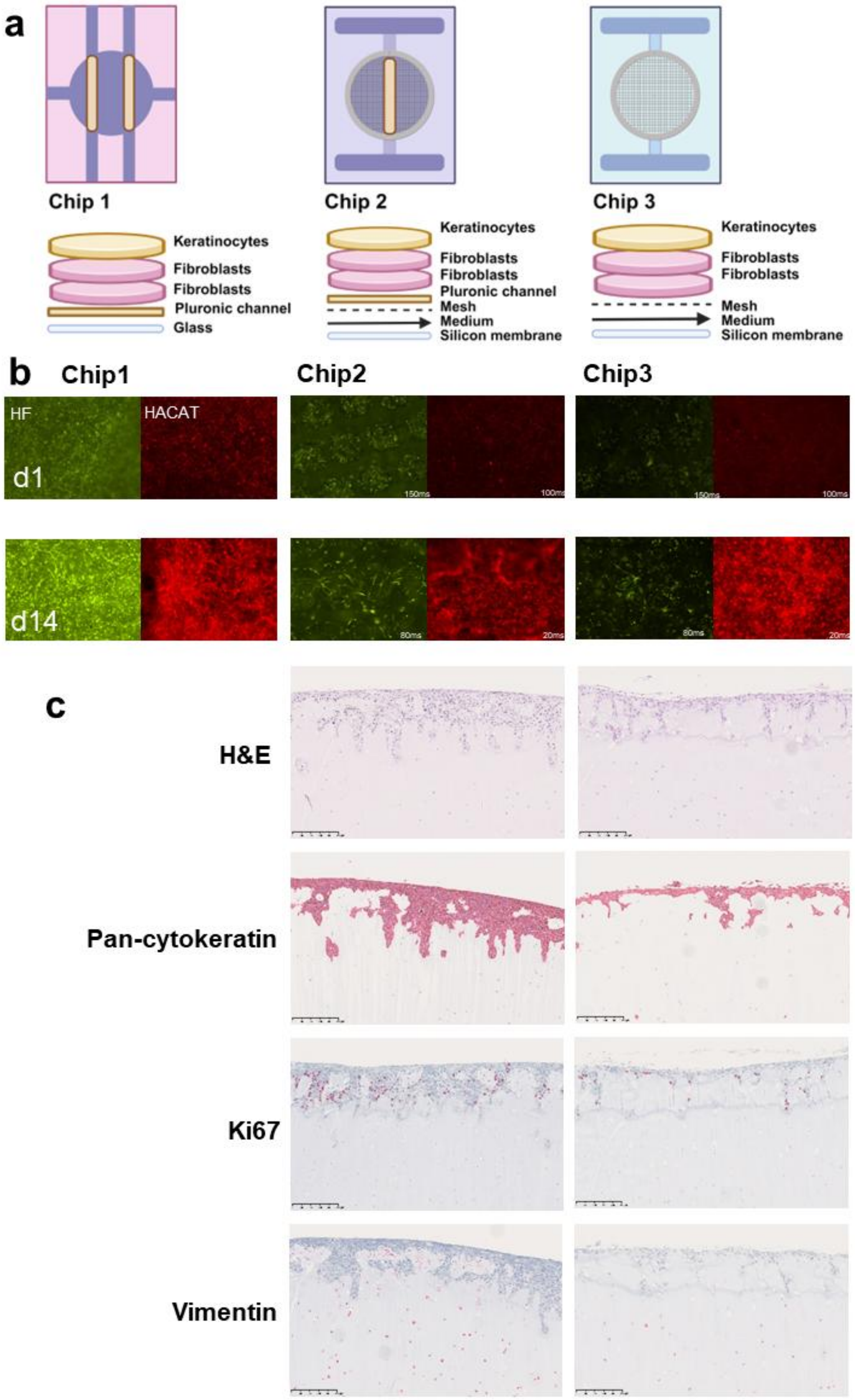


**Supplementary Figure S2. Comparison of alternative microfluidic chip designs.** (a) Schematic overview of the three microfluidic chip designs evaluated. Chip 1: laser-manufactured PMMA chip with two straight central medium channels, layered with keratinocytes, fibroblasts, a plasma channel and a glass bottom. Chip 2: stereolithography (SLA) printed resin chip with keratinocytes, fibroblasts, one central Pluronic medium channel, an additional medium chamber underneath the grid and a silicone membrane. Chip 3: Chip 2 but without a Pluronic channel. (b) Fluorescence microscopy of fibroblasts HDF EYFP (green) and keratinocytes HaCaT mCherry (red) cultured within Chip 1, Chip 2, and Chip 3 at d1 and d14 of culture. 10x magnification. (c) Histological characterization of the 3D bioprinted dermis after 2 weeks of culture. Sections were stained with hematoxylin and eosin (H&E), pan-cytokeratin, Ki67, and vimentin. Scale bar, 250 µm; 10x magnification.

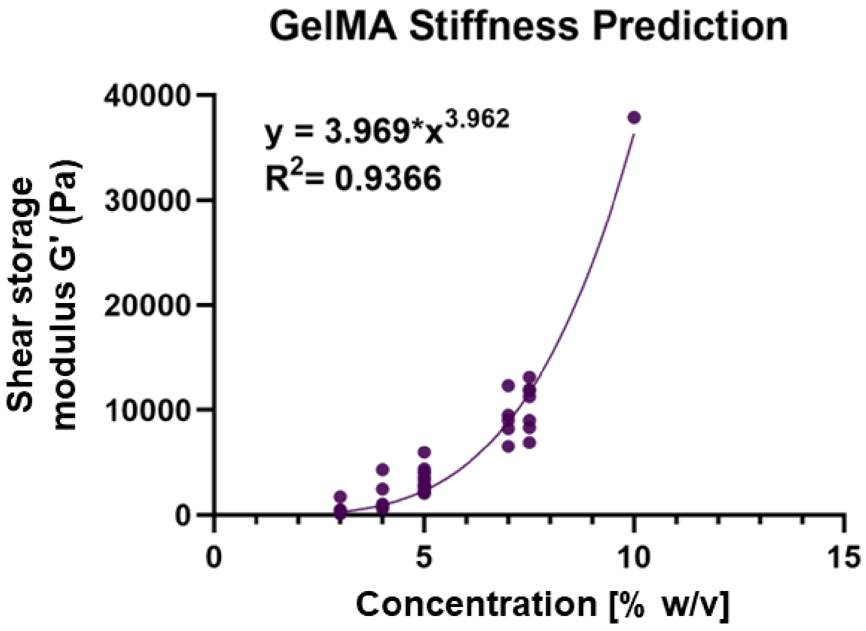


**Supplementary Figure S3. Effect of GelMA concentration on gel stiffness.** The stiffness of GelMA samples (n = 40) was measured at different concentrations (% w/v) using an ElastoSens Bio rheometer. Gel stiffness (shear storage modulus G') is shown as storage modulus G' in pascals (Pa). For each concentration, at least 5 replicates were measured, except for the highest concentration to save valuable material. Using GraphPad Prism version 11.0.0 a non-linear fit using the power series $y = a*x^b$ was performed which resulted in the formula $y = 3.969*x^{3.962}$ with an $R^2$ of 0.9366. Quality control of our GelMA batches involves routine measurement of stiffness at 5 % (w/v). This information in addition to the GelMA formula was used in subsequent experiments to estimate the necessary concentration for a defined stiffness of interest.

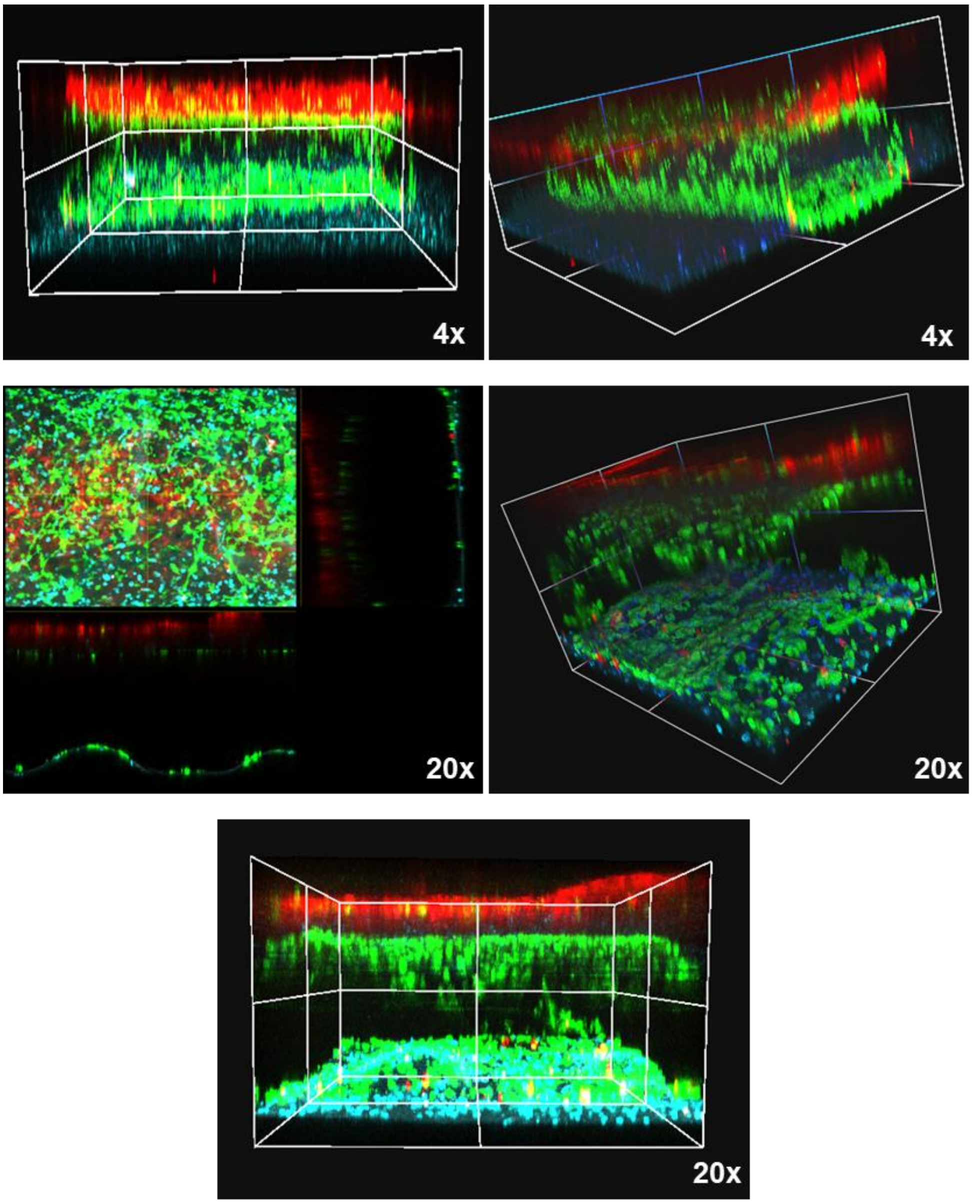


**Supplementary Figure S4. Vascularized skin-on-chip.** Dissolution of extrusion-printed, thermo-reversible Pluronic F127 ink forms open channels through the tissue. After printing, channels in the skin-on-chip were flushed with collagen solution and incubated on a custom-designed orbital shaker for 30 min at 37°C [1]. After d5 of incubation at ALI, confocal imaging was performed to visualize keratinocytes (red), endothelial cells (green) and surrounding fibroblasts (cyan). Top images show 3D reconstructed multicolor stacks acquired with a 4x objective, whereas lower images zoomed in and were acquired with a 20x objective in an FV4000 confocal microscope (Evident, Japan).

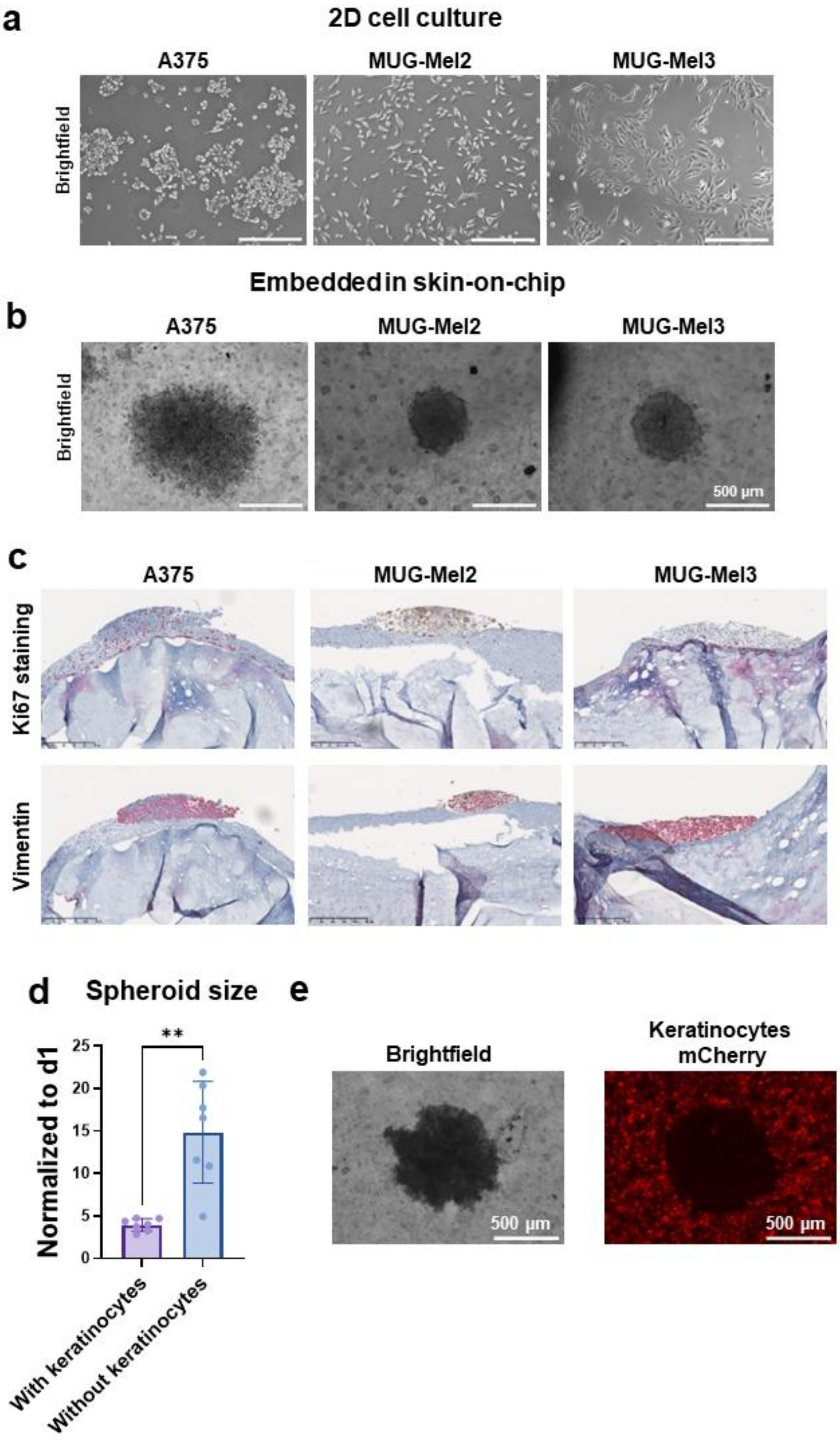


**Supplementary Figure S5. Comparison of different melanoma cell lines in 2D and 3D.** (a) Brightfield microscopy images of three melanoma cell lines (A375, MUG-Mel2, and MUG-Mel3) cultured under standard 2D monolayer conditions. (b) Comparison of A375, MUG-Mel2, and MUG-Mel3 upon integration into the skin-equivalent construct after V-bottom plate spheroid formation. (c) Immunofluorescence staining of melanoma-on-chip generated from A375, MUG-Mel2, and MUG-Mel3 cells after three weeks in culture. Ki67 staining (top row) reveals proliferative activity, while vimentin staining (bottom row) marks mesenchymal characteristics. (d) Area of spheroid on d21 normalized to d1, with and without a keratinocyte layer. (e) Fluorescence microscopy of mCherry-expressing HaCaT keratinocytes with a central melanoma spheroid protruding through the keratinocyte layer. Scale bar 500 µm.

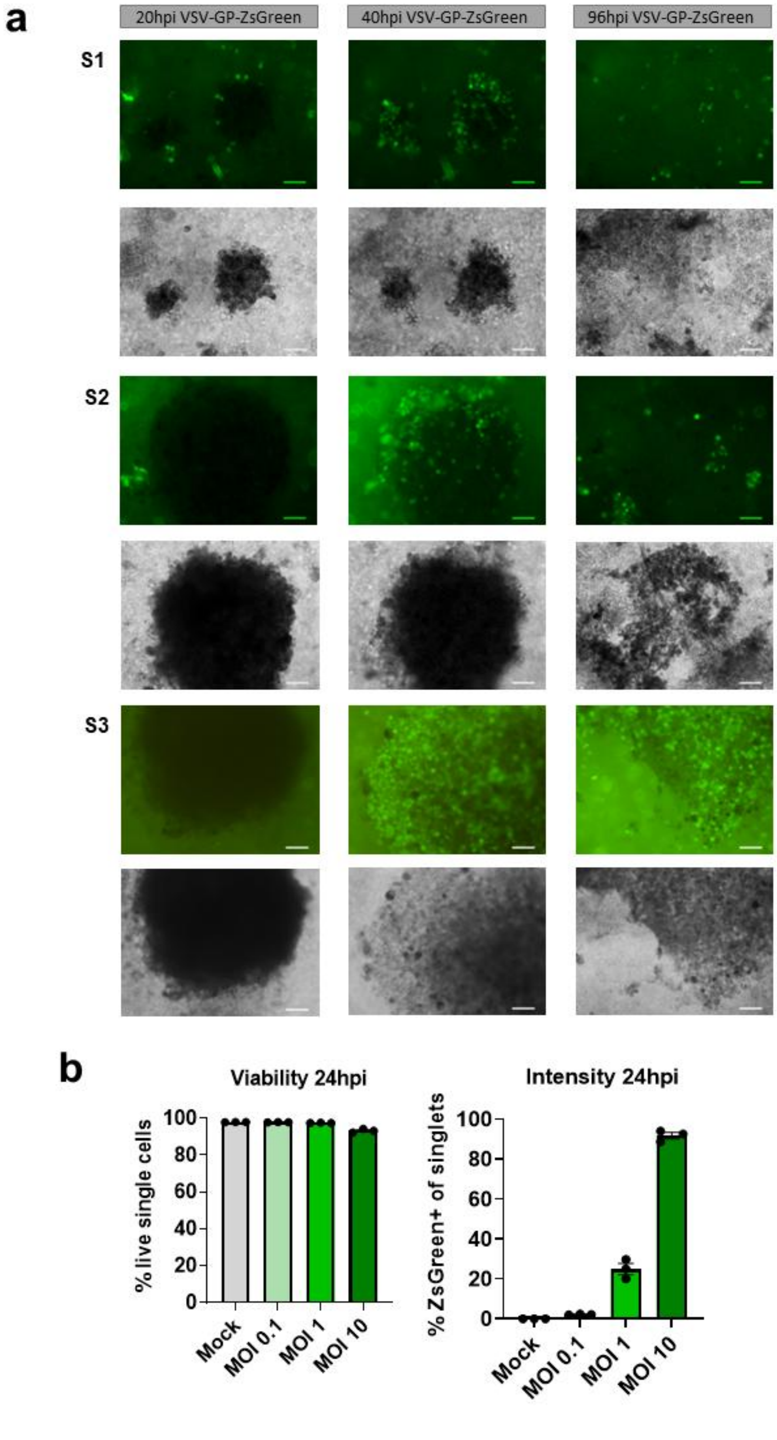


**Supplementary Figure S6. Representative VSV-GP-ZsGreen infected melanoma-on-chip.** (a) Representative GFP (top) and corresponding brightfield (bottom) microscopy images of three independent melanoma-on-chip replicates from three independent experiments (S1–S3) at 20, 48, and 96 h post-infection (hpi) with VSV-GP-ZsGreen, showing progressive spread of infection and spheroid disintegration over time. (b) Flow cytometric quantification in MUG-Mel2 cells cultured in 2D at 24 hpi, comparing mock and VSV-GP-ZsGreen (MOI 0.1, MOI 1, MOI 10) conditions: percentage of ZsGreen-positive single cells (left) and percentage of live single cells (right), n=3.

References:

1. D. Nothdurfter, C. Ploner, D. C. Coraça-Huber, et al., "3D Bioprinted, Vascularized Neuroblastoma Tumor Environment in Fluidic Chip Devices for Precision Medicine Drug Testing," Biofabrication 14, no. 3 (2022): 035002, https://doi.org/10.1088/1758-5090/ac5fb7.
